# Indexing All Life’s Known Biological Sequences

**DOI:** 10.1101/2020.10.01.322164

**Authors:** Mikhail Karasikov, Harun Mustafa, Daniel Danciu, Marc Zimmermann, Christopher Barber, Gunnar Rätsch, André Kahles

**Affiliations:** Biomedical Informatics Group, Department of Computer Science, ETH Zurich, Zurich, Switzerland; Biomedical Informatics Research, University Hospital Zurich, Zurich, Switzerland; Swiss Institute of Bioinformatics, Zurich, Switzerland; Associate faculty in the Department of Biology at ETH Zurich, Zurich, Switzerland; ETH AI Center, ETH Zurich, Zurich, Switzerland

## Abstract

The amount of biological sequencing data available in public repositories is growing exponentially, forming an invaluable biomedical research resource. Yet, making it full-text searchable and easily accessible to researchers in life and data science is an unsolved problem. In this work, we take advantage of recently developed, very efficient data structures and algorithms for representing sequence sets. We make Petabases of DNA sequences across all clades of life, including viruses, bacteria, fungi, plants, animals, and humans, fully searchable. Our indexes are freely available to the research community. This highly compressed representation of the input sequences (up to 5800*×*) fits on a single consumer hard drive (*≈*100 USD), making this valuable resource cost-effective to use and easily transportable. We present the underlying methodological framework, called MetaGraph, that allows us to scalably index very large sets of DNA or protein sequences using annotated De Bruijn graphs. We demonstrate the feasibility of indexing the full extent of existing sequencing data and present new approaches for efficient and cost-effective full-text search at an on-demand cost of $0.10 per queried Mpb. We explore several practical use cases to mine existing archives for interesting associations and demonstrate the utility of our indexes for integrative analyses.

## Introduction

For more than a decade, continuing innovation in the area of high-throughput sequencing has propelled research in the biomedical domain and led to an exponential growth in worldwide sequencing capacity^1^. The number of sequenced nucleotides contained in the European Nucleotide Archive (ENA) doubles roughly every 45 months and currently makes up over 5.5 *·* 10^16^ nucleotide bases (55 Petabases)^2^ of raw read data. The currently supported, classical pattern for accessing sequence data in public repositories is to identify relevant records using descriptive metadata and to retrieve a copy or a slice of the data for further processing. This either involves downloading the records or accessing them on a cloud platform – both approaches often require significant resources. However, the raw sequencing data itself remains inaccessible for full-text search, significantly limiting its potential for future research. To address this issue, we propose a highly scalable approach to index Petabase-size repositories of raw sequencing data, transforming them into a highly compressed representation that is more accessible for downstream analysis. Inspired by recent advances in sequence algorithms and data structures^3–10^, we focused on specific technical developments for higher compression and faster search^11,12^ to demonstrate that indexing the repositories like the Sequence Read Archive (SRA) as a whole is not only theoretically possible but practically feasible.

Indexing sequence data at such a scale gives rise to a variety of technical challenges for which some solutions have been proposed in the recent past. Naturally, one focus is on making the genetic variation in large cohorts, especially in human, accessible for research and clinical use. Only very recently, frameworks for variation graphs, such as VG^13^, and methods for compressing haplotypes^14^, or paths in graphs more generally^15^, have improved variation-aware alignment and variant calling in general^16,17^. However, these methods show strong limitations in the scalability of input size and variability. A second focus of algorithmic work is on taxonomic classification and read annotation, where the task is to match a given query sequence against a (large) set of known reference sequences or taxonomic labels. The classic approach for alignments against a collection of assembled genomes is BLAST^18^. Although the tool is heavily used and has been gradually improved for speed^19,20^ over the past 30 years, it still lacks the scalability that is required for high-throughput searches on highly diverse sequence collections. With approaches like Centrifuge^21^ promising speed-up has at least been gained for jointly querying large sets of assembled genomes. A third technical focus is on the *sequence* or *experiment discovery problem*: querying a sequence of interest (e.g., a transcript) against all samples available in a sequence repository. Current approaches for experiment discovery can be grouped into three main categories: (i) methods based on sketching techniques, which summarize the input data using one or multiple hashing operations and then use these summaries (sketches) to estimate distances between the query and a target^22–24^; (ii) methods employing Bloom filter-based data structures to allow for approximate membership queries^25–29^; (iii) methods for the exact representation of *annotated De Bruijn graphs*, also called *colored De Bruijn graph*, storing additional metadata as annotations of its nodes or edges^4^, such as Mantis^6,30^ and others^31–33^. All methods approach experiment discovery by matching short substrings of a fixed length *k*, called *k-mers*, from query sequences against those stored in the index. While efficient, they often lack the ability for sensitive alignment ^34^. In addition to the above work, there has also been growing interest in the use of De Bruijn graph-based indexes for alignment tasks as a way to accelerate alignment to repeat-prone reference genomes^35^ or to pangenomes and unassembled read sets^5,36–38^.

A major challenge faced by all discussed methods is to unite the ability to operate efficiently on Petabase-scale sequence collections while supporting fast and versatile query operations, such as search, alignment, and experiment discovery. To bridge this evident gap and to demonstrate the practical feasibility of full-text indexing the whole SRA, we present MetaGraph, a versatile framework for the indexing and analysis of biological sequence libraries at petabase-scale.

## Results

### Efficient, modular, and extensible sequence representation with MetaGraph

Building on recent advances in the field and by developing additional approaches for sequence analysis, we devised the MetaGraph framework that forms the core of this work and scales from use on single desktop computers up to distributed compute clusters. Notably, MetaGraph can index biological sequences of all kinds, such as raw DNA/RNA sequencing reads, assembled genomes, and protein sequences.

The *MetaGraph index* consists of an annotated sequence graph (**Figure 1, right top**) that has two main components: The first is a *k-mer dictionary* representing a De Bruijn graph (**Figure 1, right middle**). The *k*-mers stored in this dictionary serve as elementary tokens in all operations on the MetaGraph index. The second component is a representation of the metadata encoded as a *relation* between *k*-mers and any categorical features (called *annotation labels*), such as sample IDs, geographic locations^39^, quantitative or positional information. This relation is represented as a sparse matrix (*annotation matrix*; **Figure 1, right middle**). For instance, for binary annotation, the (*i, j*)^th^ element of this matrix has value 1 if and only if the *i*^th^ *k*-mer is associated with the *j*^th^ feature. We employ various techniques to represent the De Bruijn graph and the annotation matrix in a highly compressed form^3,11,12,40^. We describe these techniques in **Methods**. The MetaGraph framework supports using different graph and annotation representations interchangeably, adapting to different storage requirements and analysis tasks, and allowing for easy adoption of the latest algorithmic developments. Consequently, we made certain design choices when developing MetaGraph: (i) Use of succinct data structures and efficient representation schemes for extremely high scalability; (ii) Efficient algorithmic use of succinct data structures (e.g., prefering batched operations); (iii) Modular architecture supporting several graph and annotation representations and enabling the *addition* of new algorithms with *little code overhead*. – These strengths allow for MetaGraph to easily adapt to the rapid methodological advancements for sequence set representation^41–44^.

**Figure 1:**
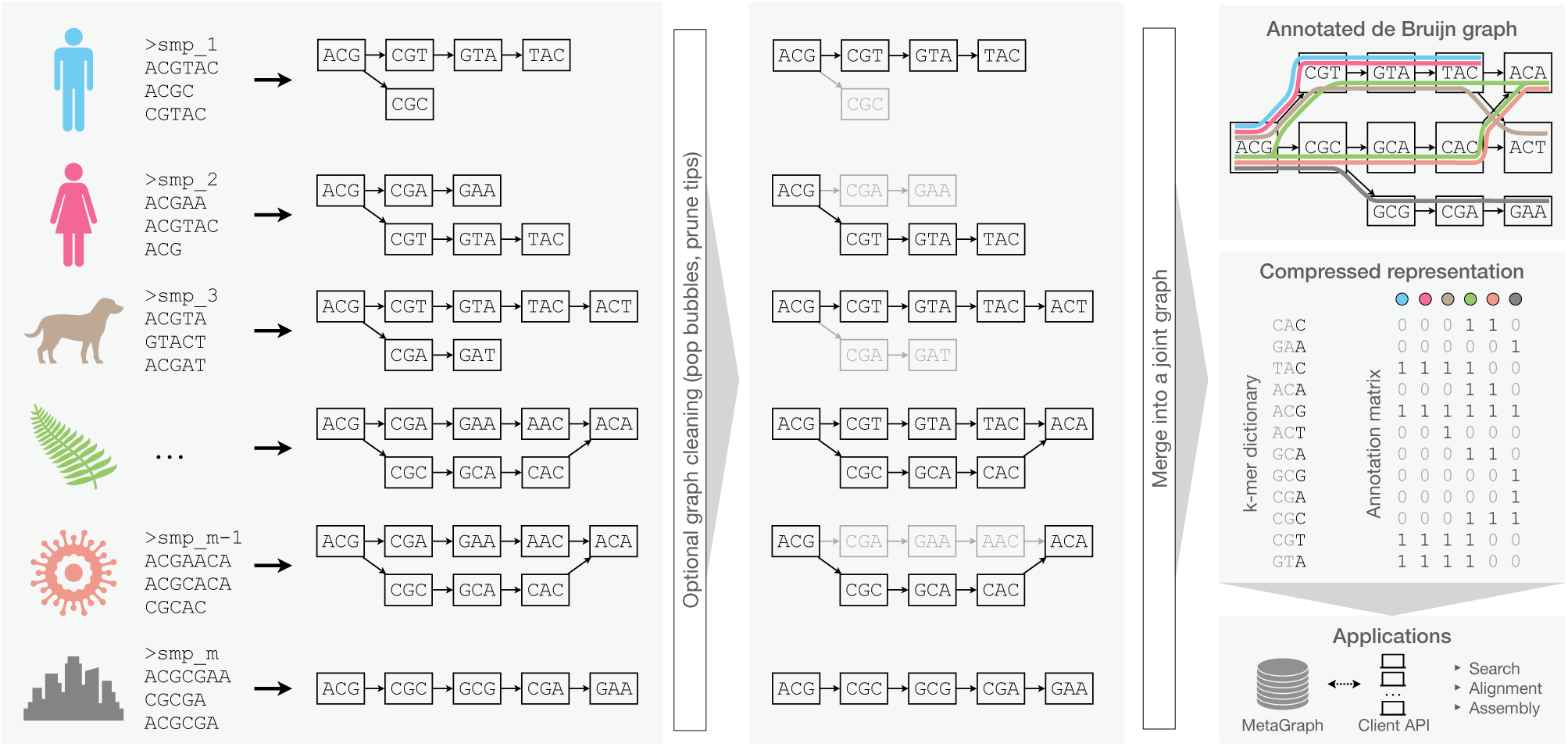
The MetaGraph framework. Schematic overview of graph construction and representation. Individual sequencing samples (**left**) are assembled into graphs (*sample graphs*), which are then cleaned to remove spurious paths and misassemblies (**middle**). Next, the sample graphs are merged to form a MetaGraph index (**right top**), consisting of the compressed *k*-mer dictionary index and a compressed annotation matrix (**right middle**, gray color indicates parts that can be efficiently compressed and do not have to be stored explicitly). The index is then used as the basis for downstream applications, such as sequence search, differential assembly, and other queries (**right bottom**).

### A workflow for scalable multi-sample index construction

Without loss of generality, we will assume that the indexing is applied to read sets that stem from sequencing biological samples, where each annotation label encodes a sample ID. Thus, the annotation matrix encodes the samples to which each *k*-mer belongs. Our indexing workflow proceeds in three steps: (i) data preprocessing, (ii) graph construction, and (iii) annotation construction. (i) Data preprocessing involves the construction of separate De Bruijn graphs (*sample graphs*) from the raw input samples (**Figure 1, left**). We optionally apply a subsequent cleaning step to each sample graph to reduce the impact of potential sequencing errors (**Figure 1, middle**). (ii) In graph construction, all sample graphs obtained in the first stage are merged into a single joint De Bruijn graph (**Figure 1, right top**). (iii) In annotation construction, we build the columns of the annotation matrix to indicate the membership of different *k*-mers to their respective sample graphs (**Figure 1, right middle**). Finally, the graph and the annotation are compressed into representations best suited for their target application (**Methods**).

### First-in-class scalability of MetaGraph

We evaluated MetaGraph’s scalability against other state-of-the-art indexing tools: BIGSI^25^, COBS^26^, Bifrost^45^, and Mantis^30a^, on subsets of increasing size up to 25,000 read sets randomly drawn from the entire collection of bacterial and viral genomic read sets^25^. While all the representations grow in size approximately linearly with the input size, the absolute values are drastically different (**Figure 2 a**). The space taken by MetaGraph indexes in the smallest representation is 16–38 times smaller than that of the other evaluated approaches, despite some of them using a lossy compression approach (BIGSI and COBS).

**Figure 2:**
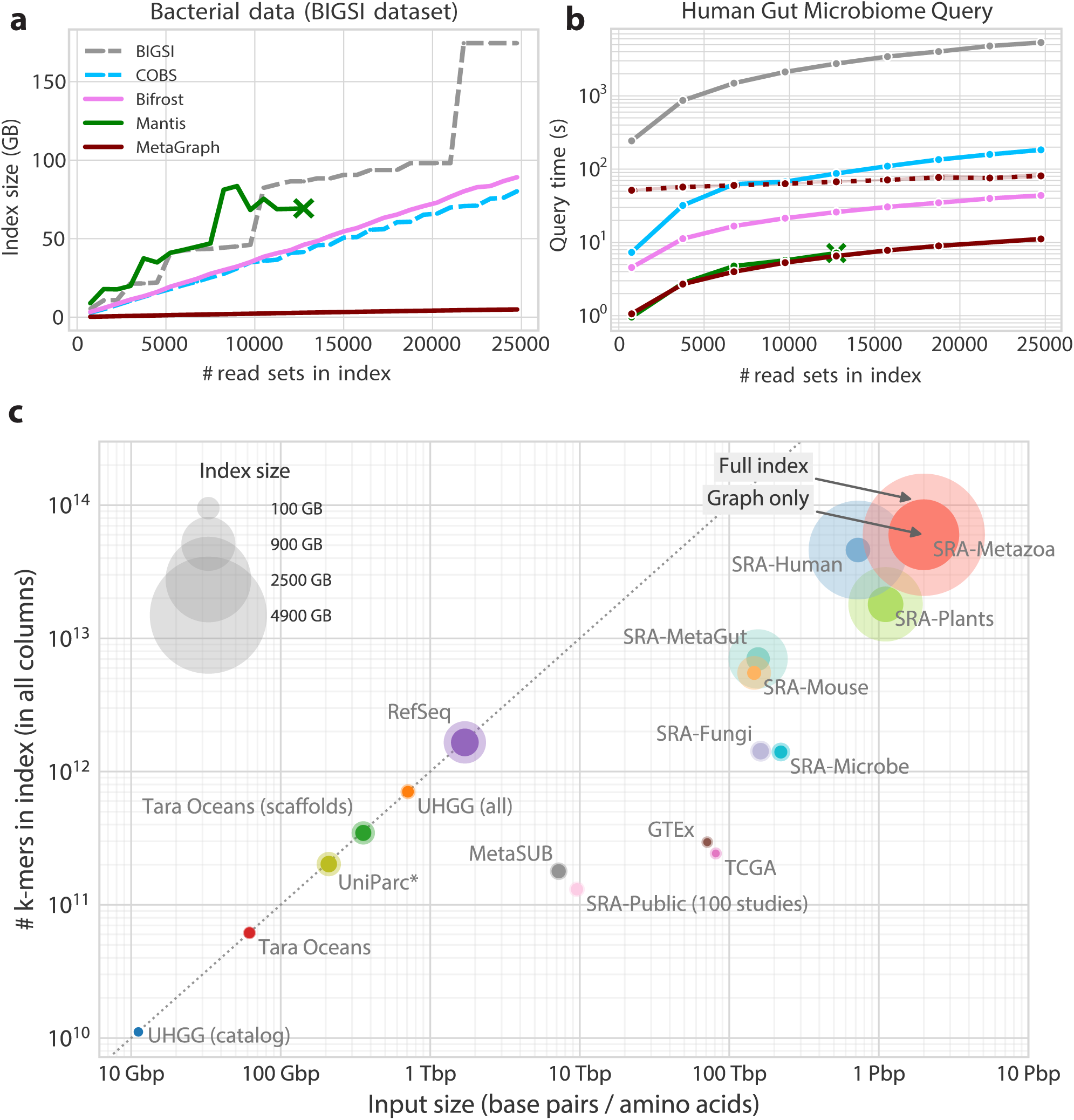
**a)** Size of evaluated index data structures for representing a set of microbial WGS experiments of increasing size, shown for both lossy indexing methods BIGSI and COBS and lossless Mantis, Bifrost, and MetaGraph with the SuccinctDBG and RowDiff-MultiBRWT compression schemes to encode the graph and the annotation, respectively. Dashed lines indicate lossy methods. The construction of the Mantis indexes could not be done on all the subsets as indexing of the 13,500 sample subset was only 43.6% complete after reaching our 240 h run time limit. **b)** Times for querying human gut metagenome AMPLICON sequencing reads (SRA accession ID `DRR067889`) against indexes constructed with MetaGraph and other state-of-the-art tools from sets of microbial WGS experiments of increasing size. All curves show the performance of exact k-mer matching, except for the dotted MetaGraph curve, which shows the query performance with the more sensitive search strategy involving alignment. **c)** Overview of all MetaGraph indexes — For all data sets we show the total number of input characters on the x-axis and index size (given as the total number of unique *k*-mers) on the y-axis. Marker size represents the size of the index. The solid portion of each marker represents the fraction of the total size taken by the graph and the translucent portion represents the fraction taken by the annotation (**Table 1**). (*) Input of the UniParc dataset are amino acid sequences, not base pairs.

**Table 1:** Summary of the data sets and constructed indexes. Indexes marked with ^†^ encode genome coordinates (hence, fully lossless). Indexes marked with ^‡^ in addition to the membership of *k*-mers to different labels also represent their respective counts. Each index is assigned to one of three classes: DNA (D), RNA (R), or Protein (P). The index sizes for SRA-Metazoa and SRA-Public marked with * are estimated from their respective subsets. The compression ratio is measured in char/byte (base pairs or amino acids of input sequences per byte of the MetaGraph index), which corresponds to the input/output ratio if 1 char. takes 1 byte. The last two columns show the two factors of the compression ratio: redundancy of the data (due to sequencing depth, sequencing errors, and genome complexity/repetitiveness), and indexing efficiency, i.e., how many (*k*-mer, label) relations are encoded per byte of the MetaGraph index.

| Type | Data set | Class | Tbp | # labels | Index size | Compression (char./byte) | Redund. (char./rel.) | Efficiency (rel./byte) |
| --- | --- | --- | --- | --- | --- | --- | --- | --- |
| Assembled seq. | UHGG (catalog) | D | 0.01 | 4,644 | 3.2 GB | 3.5 | 1.0 | 3.5 |
|  | UHGG (all) | D | 0.71 | 286,997 | 27.3 GB | 26.0 | 1.0 | 25.9 |
|  | Tara Oceans (scaffolds) | D | 0.36 | 318,205,057 | 110.2 GB | 3.2 | 1.0 | 3.1 |
|  | Tara Oceans with coord.<sup>†</sup> | D | 0.06 | 34,815 | 14.6 GB | 4.2 | 1.0 | 4.2 |
|  | RefSeq with coordinates<sup>†</sup> | D | 1.70 | 85,375 | 508.9 GB | 3.3 | 1.0 | 3.3 |
|  | UniParc | P | 0.21 | 543,904,874 | 124.6 GB | 1.7 | 1.0 | 1.6 |
| RNA-Seq | GTEX | R | 71.2 | 9,759 | 9.6 GB | 7,416 | 241.2 | 30.7 |
|  | GTEX with counts<sup>‡</sup> | R | 71.2 | 9,759 | 76.3 GB | 934 | 241.2 | 3.9 |
|  | TCGA | R | 81.2 | 11,095 | 11.1 GB | 7,288 | 334.4 | 21.8 |
|  | TCGA with counts<sup>‡</sup> | R | 81.2 | 11,095 | 81.2 GB | 1,001 | 320.4 | 3.1 |
| MGS | MetaSUB | D | 7.2 | 4,220 | 46.7 GB | 155 | 40.5 | 3.8 |
|  | SRA-MetaGut | D | 155.8 | 241,384 | 1,111.3 GB | 140 | 22.2 | 6.3 |
| SRA subsets | SRA-Microbe | D | 221.1 | 446,506 | 57.1 GB | 3,870 | 157.6 | 24.5 |
|  | SRA-Fungi | D | 162.1 | 121,900 | 80.5 GB | 2,013 | 113.9 | 17.7 |
|  | SRA-Plants | D | 1,109.2 | 531,714 | 1,844.1 GB | 602 | 61.5 | 9.8 |
|  | SRA-Metazoa (Human) | D | 725.4 | 436,494 | 3,402.1 GB | 213 | 15.7 | 13.5 |
|  | SRA-Metazoa (Mouse) | D | 146.6 | 57,938 | 291.6 GB | 503 | 26.6 | 18.9 |
|  | SRA-Metazoa (w/o Human) | D | 1,999.5 | 805,239 | 5,366.5 GB | 373 | 33.3 | 11.2 |
| SRA | SRA-Public (100 studies) | D | 9.6 | 5,184 | 32.0 GB | 300 | 73.3 | 4.1 |
|  | SRA-Public to 01.01.2023 | D | 38,949.9 | 23,010,648 | 130 TB* | 300* | 73.3* | 4.1* |

However, this exceptional space-efficiency of MetaGraph does not compromise query time. We devised several efficient search algorithms to identify matching paths in the De Bruijn graph and retrieve the corresponding annotations. Briefly, for *k-mer query*, input sequences are broken down into *k*-mers that are simply mapped to the nodes of the graph. Then, the respective annotations are retrieved and the aggregated result is returned as output (**Extended Data Figure 1 c**). For increased sensitivity, we developed algorithms for *sequence-to-graph alignment*^40,46^, which identify the closest matching path in the whole graph (**Extended Data Figures 1 d; Methods**). We also designed a batch query algorithm (schematic in **Extended Data Figure 1 e, Methods**), exploiting the presence of *k*-mers shared between individual queries by forming a fast intermediate *query subgraph*, that increases throughput up to 100-fold for large repetitive queries (e.g., sets of sequencing reads).

To evaluate the performance of the sequence search strategies implemented in MetaGraph against the existing methods for indexing raw sequencing data, we queried four different types of sequences: a set of antimicrobial resistance (AMR) gene DNA sequences from the CARD database^47^, reads from an environmental metagenome sample ^b^ from MetaSUB^48^, AMPLICON reads from a human gut metagenome sample (`DRR067889`), and metagenomic WGS reads (`SRR322873`). We find that MetaGraph is highly competitive in query time compared to all other evaluated methods (**Figure 2 b**), while maintaining orders of magnitude smaller index representations. Moreover, using batch query optimizations (**Methods**) MetaGraph shows the best performance on sets of highly redundant queries (`DRR067889` and `SRR322873`). Even the sensitive search employing sequence alignment in MetaGraph is considerably faster than BIGSI, a tool for approximate k-mer matching originally developed to index this data set. In general, the retrieval of *k*-mer labels based on exact *k*-mer matching can be seen as an approximation of semi-global alignment, where the query sequence would otherwise be aligned against the subgraphs induced by those annotation labels^6,25,26^. If each annotation column is constructed from a single genome, this process mirrors the classical problem of alignment against a collection of reference genomes. This approximation becomes less accurate as subgraph complexity increases (**Extended Data Figure 2**).

### A Petabase-scale index for public sequence data

As we set out to demonstrate the practical feasibility of indexing entire sequence archives, we applied MetaGraph to index a substantial part of the public section of the NCBI Sequence Read Archive (SRA), including both DNA and RNA, alongside restricted-access human cohorts and amino acid sequences (present in the UniParc database). Consequently, our indexed data sets range from large RNA-Seq cohorts (TCGA^49^, GTEx^50^) and vast archives of genome sequencing records from the NCBI Sequence Read Archive (SRA)^51^ (comprising 2,347,037 samples of microbial, fungal, plant, and metazoa organisms) to large sets of highly diverse whole metagenome sequencing samples (MetaSUB^48^ cohort and all publicly available human gut metagenome samples from SRA) and collections of reference and assembled sequences (RefSeq^52^, UHGG^53^, and Tara Oceans^54^). Our MetaGraph index of the complete UniParc dataset^55^ demonstrates the straightforward use of protein sequences as input data. For selected data sets, we generated indexes preserving *k*-mer counts or coordinates utilizing the concept of *counting De Bruijn graphs*^40^. The key statistics of all data sets presented in this work and MetaGraph indexes constructed from them are provided in **Table 1** and visualized in **Figure 2 c**. The resulting indexes form a valuable community resource, as they succinctly summarise large sequence data sets while supporting a variety of sequence queries against them.

When processing raw sequencing data, we applied moderate cleaning on the inputs (**Methods**), which led to significant reductions in the required storage space without a significant effect on index completeness and query sensitivity. In total, we processed 4.8 Petabases (*≈* 2.5 Petabytes) of input sequences, which finally makes this data fully and efficiently searchable by sequence.

### Data redundancy drives compression ratio

MetaGraph’s compression performance greatly depends on the properties of the sequence sets being indexed. Since the input data comes in different formats (FASTA, FASTQ, SRA, etc.), we measured the final compression ratio in characters/byte (i.e., the average number of base pairs or amino acids of input per byte of the MetaGraph index) to make it comparable across different data sets. For a more detailed analysis, we decomposed the compression ratio into two factors: the *data redundancy* and the *indexing efficiency* shown in the last two columns of **Table 1**, respectively. Intuitively, the redundancy shows the amount of data duplication *within an indexed sample*, while the indexing efficiency reflects data duplication *across different samples* and increases for sets of related samples.

While the GTEx ^50^ and TCGA ^49^ cohorts amount to over 100 TB of raw compressed RNA sequencing data, they show only limited diversity and our MetaGraph index represents both in *≈* 10 GB space, thereby achieving the highest compression ratios among all the data sets we have indexed (up to 7,416 bp/byte). Even when adding *k*-mer counts, the final compression ratio is about 1,000 bp/byte.

The medium complexity range mainly consists of whole genome sequencing read sets, usually showing more diversity and lower coverage than the human transcriptome cohorts and, thus, showing less redundancy. At the time of collection (April 2020), the SRA data sets we indexed comprised over 90% of all publicly available samples in the selected groups. Only for the SRA-Microbe data set, we relied on a previously defined set of 446,506 virus, Prokaryote, and small Eukaryote genomes used in the original BIGSI publication^25^. Notably, the fully searchable and annotated MetaGraph representation of the SRA-Microbe data set takes only 57 GB, which is 28*×* smaller than the 1.6 TB taken by the BIGSI index^25^.

On the opposite end stand whole metagenome shotgun sequencing cohorts. Here, we have selected the MetaSUB cohort^48^, containing 4,220 environmental metagenome samples comprising 7.2 Tbp, and the SRA-MetaGut cohort, containing all human gut metagenome samples available on SRA at this time, comprising approximately 156 Tbp. These inputs represent very diverse populations of organisms and contain many rare sequences. Consequently, compression ratios are lower (140–155 bp/byte) but still large enough to compactly represent those vast collections in searchable indexes (46.7 GB for MetaSUB; 1,111 GB for SRA-MetaGut).

Lastly, collections of assembled genomes and protein sequences feature the highest diversity and the least data redundancy, with the similarity between samples being a function of evolutionary distance. Even though the lack of inherent redundancy within these data sets makes them hard to compress, MetaGraph allows indexing them to make them fully searchable while keeping the index small.

The complete lists of sample identifiers used in our experiments are provided in Supplemental Data.

### Indexes are highly-accurate and near-complete

As our workflow for indexing raw read sets involves per-sample cleaning procedures, in the context of *experiment discovery*, a given query sequence may not retrieve a true label because of missing *k*-mers that were filtered out. In this light, we evaluated the accuracy of experiment discovery on our SRA-derived MetaGraph indexes. We validated our cleaning and sequence-to-graph alignment approach by showing that each of the individual sample graphs is an accurate representation of their respective input read sets, with *∼*82% of our query reads realigning back to their respective sample graphs with at least 80% sequence identity (**Extended Data Figure 3 a**). Notably, even when mapping these reads to their corresponding annotated graph indexes with k-mer matching (without sequence alignment), we similarly observe that 80% of the query reads retrieve their true labels with at least 80% sequence identity (**Figure 3 a**).

**Figure 3:**
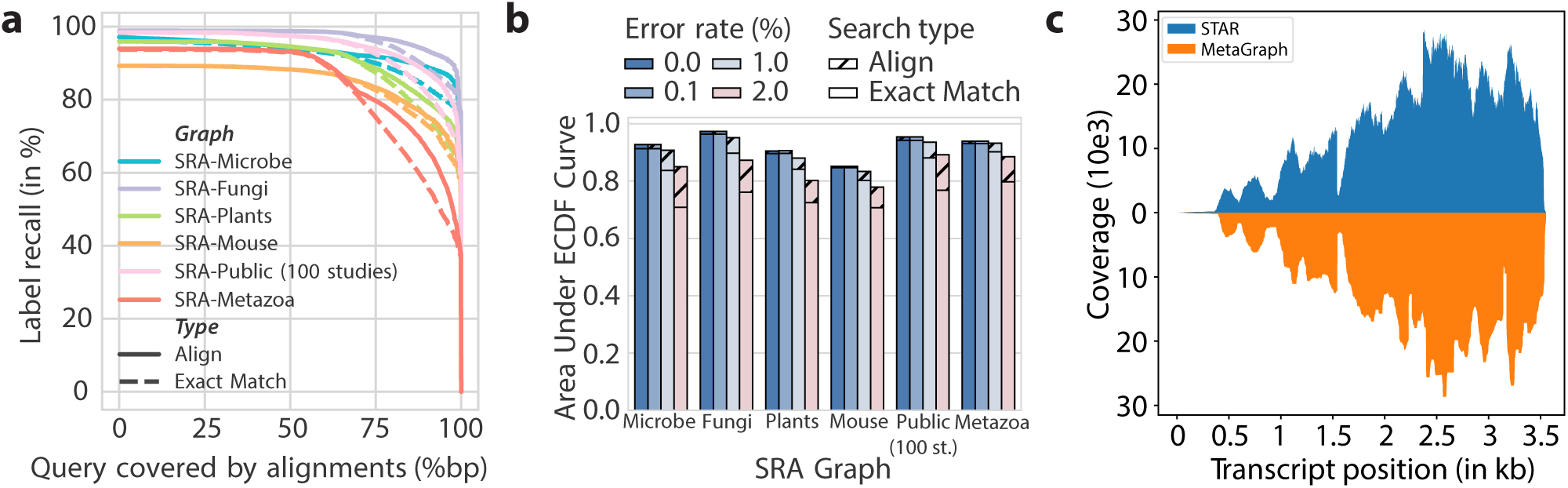
MetaGraph completeness and accuracy. **a)** Complementary estimator of the cumulative distribution function (ECDF) curve showing the fraction of reads that are retrieving the correct label from the index (*Label recall*, y-axis) requiring at least x% matching positions (x-axis) for a range of indexed SRA datasets. We define the *realignability* of a graph-query pair as the area under this curve. **b)** Realignability of each graph for increasing error rates in the query. **c)** Coverage plot for human transcript SFTPB-207 in GTEx sample SRR599154 determined via STAR alignment against hg38 (top, blue) and retrieved from the MetaGraph GTEx index with counts (bottom, orange).

In addition, we evaluated the robustness of our mapping algorithms to sequence variation using altered versions of the query reads with varying mutation rates. Generally, we observe that the experiment discovery accuracy difference between exact *k*-mer matching and sequence-to-graph alignment increases with increasing sequence variation (**Figure 3 b**). So far, we have only considered the presence or absence of an annotation label. Especially for functional sequencing data, such as RNA-Seq, it is crucial that MetaGraph can encode per-sample count information. As an example, we provide the expression of transcript SFTPB-207 (ENST00000519937.6) in a sample from the GTEx cohort (**Figure 3 c**). Even though our representation is a 1,000 fold compression of the original data, we can closely reproduce the coverage profile generated from aligning the original raw sequences to the reference using STAR. As all count information is encoded per sample, tissue specificity is preserved. For this example, we find the same surfactant transcript mainly expressed in lung, and partially in testis, but not elsewhere (**Extended Data Figure 3 b**) – as expected.

### Exploring the human gut resistome and phageome

To showcase MetaGraph’s utility in real-time -omics analysis on very large-scale data sets, we queried the full CARD antimicrobial resistance (AMR) database^47,56^ and all bacteriophages from RefSeq Release 218^57^ against the 241,384 human gut microbiome samples in our SRA-MetaGut database (**Methods**). Notably, this is an analysis that would otherwise have required access to hundreds of terabytes of raw sequencing data.

We recover strong associations between the *Escherichia λ* phage ev017 and the *E. coli β*-lactamase gene, two *Klebsiella* phages and carbapenem-resistant *β*-lactamases from *Klebsiella pneumoniae*, and *λ* ev017 and the RNA antibiotic efflux pumps (**Figure 4 a**). In addition, we studied trends in antibiotic resistance over time across various continents. We find significant strong growth trends over time in resistance against diaminopyrimidines in Africa, antiseptics and fluoroquinolones in Oceania, and against cephamycin and one of the so-called “last-resort” antibiotics tigecycline^58^ (the only glycylcycline approved as an antibiotic^59,60^) in South America (**Figure 4 b**).

**Figure 4:**
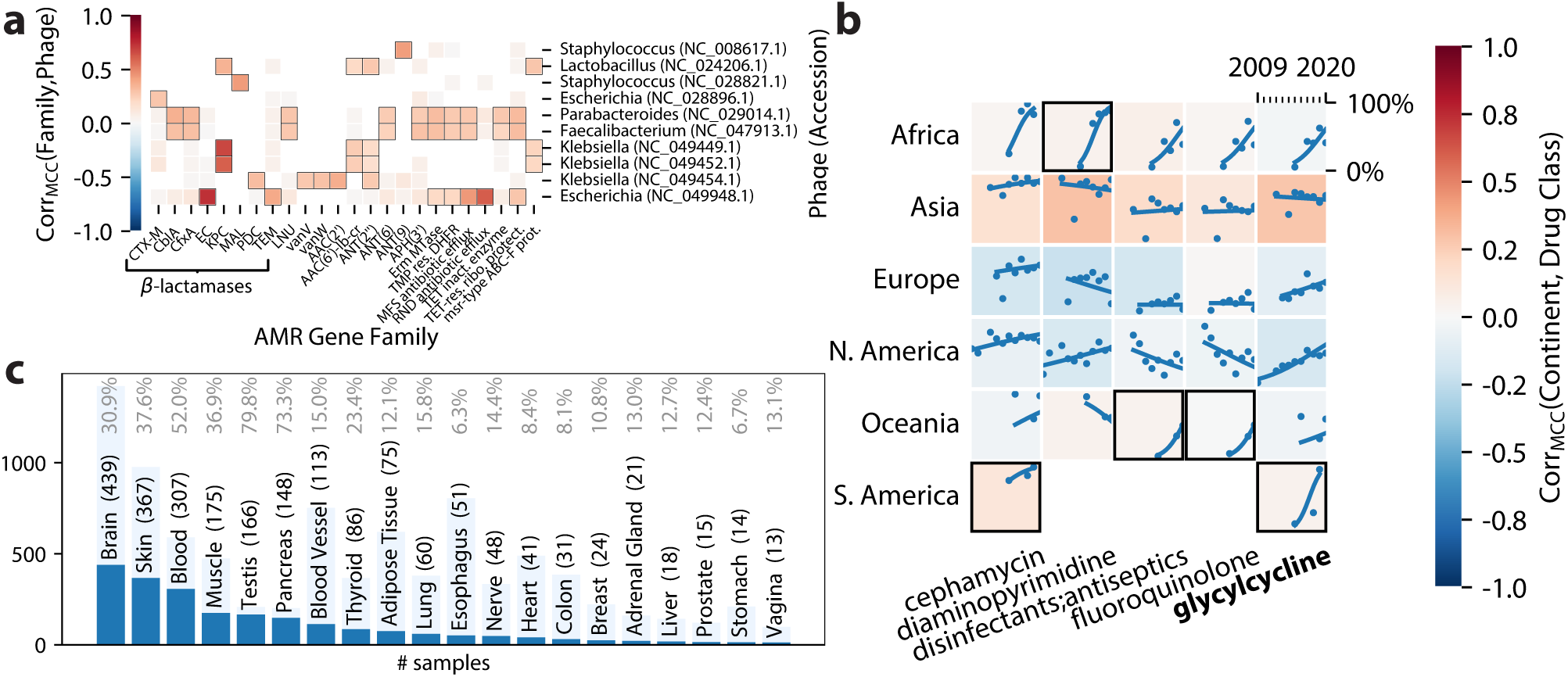
MetaGraph aids biological discovery. **a)** Associations between antimicrobial resistance (AMR) gene families and RefSeq bacteriophages (phages) discovered in publicly-available human gut microbiomes deposited to the NCBI SRA from 2008 to 2022. Squares with a black border indicate a statistically significant association and missing squares indicate hypotheses that were not tested due to an insufficient number of matching samples. **b)** Growth in the prevalence of resistance to antibiotics in human gut microbiomes across six continents from 2010 to 2020. Each point represents the normalised proportion of samples from a continent matching genes conferring resistance in a given year, while each line represents a binomial regression fit. Squares with a black border indicate regressions with significantly high training scores and missing squares indicate experiments with an insufficient number of samples. **c)** Relative fraction of GTEx samples per tissue (x-axis) that carry recurrent back-splice-junctions (BSJ). Per-tissue ratio is computed as samples that carry at least 2 BSJ (dark blue) over all samples of that tissue (light blue) - both on the y-axis. Absolute numbers are given in parenthesis, and relative fractions are in grey at the top.

As a demonstration for MetaGraph’s interactive capabilities, we provide publicly available example scripts^c^, querying the full CARD database against an index containing 4,220 whole metagenome sequencing samples from the MetaSUB cohort. We reproduce a ranking of cities based on the average number of AMR-markers in a sample (**Extended Data Figure 4 b**), consistent with the analysis performed on the raw data using orthogonal strategies^39^. We also show other exploratory analyses, linking sample metadata, such as surface material at the sampling location, to the query results (**Extended Data Figure 4 c**).

### Survey of back-splice junctions in GTEx and TCGA

Being constructed from raw data, MetaGraph indexes can easily express transcriptome features that are hard to represent in a classial, linear coordinate system. One example are back-splice junctions (BSJ) that connect the donor of an exon site to the acceptor site of a preceding exon, forming circular RNAs, which have been found to commonly occur also in humans^61^. When systematically querying all BSJ that can possibly be created using the GENCODE annotation (version 38) and the hg38 reference (**Methods**), we found a total of 672 candidates perfectly matching to the full GTEx graph that did not match to the reference genome or transcriptome, and mapped more than 90% of *k*-mers to at least one sample. While such features are hard to map using linear RNA-Seq aligners (**Extended Data Figure 4 d**), it is easy using a graph index. After removing 39 outlier-BSJ candidates that mapped to more than 300 samples, we found BSJ to be recurrently detectable in a large fraction of the GTEx tissues (**Figure 4 c**), where pancreas, testis, skin, and muscle tissues showed the largest relative fraction.

### Seamless distribution and interactive use

In addition to the single-machine use case, where the index is constructed and queried locally, MetaGraph also supports querying indexes provided on a remote server via a client-server architecture. In this approach, a set of graphs and annotations can be easily distributed across multiple machines. Each machine runs MetaGraph in *server mode*, hosting one or multiple indexes and awaiting queries on a pre-defined port (**Extended Data Figure 1 b**). This setup makes it straightforward to execute user queries across all indexes hosted on multiple servers. For easy integration of results and coordination of different MetaGraph instances, we provide client interfaces in Python (**Extended Data Figure 1 a**).

Notably, our distribution approach can be used not only for hosting multiple indexes of distinct sources but also when indexing a single data set of extremely large size. When indexing the SRA as a whole, the final MetaGraph index constructed from it exceeds the amount of memory available on a single machine. Hence, the entire data set can be split into multiple parts, where each part is indexed separately. The resulting MetaGraph indexes can then be independently loaded and hosted on multiple machines. This distribution approach enables virtually unlimited scalability. For search, each query is sent to all index partitions, followed by aggregation of the partial results to generate a final result of the query returned back to the user.

### Search engine for biological sequences MetaGraph Online

Building on MetaGraph’s distribution capabilities, we developed a search engine for biological sequences MetaGraph Online to make our indexes publicly available for online queries. The service is publicly available for user queries at https://metagraph.ethz.ch/search. It uses the MetaGraph framework for hosting the indexes and performing queries on them in real time, showing the result of a query practically immediately.

The search engine has a clean and intuitive graphical web user interface (UI) **Supplemental Figure S-7**, allowing the user to paste an arbitrary sequence and search it against a selected index. By default, the search is performed via basic *k*-mer matching. For greater sensitivity, it is also possible for all indexes to additionally align the searched sequence to the annotated graph. If *k*-mer coordinates or counts are represented in the queried index, the web UI allows retrieving them for the query sequence too.

MetaGraph Online currently hosts a large subset of the various indexes constructed and presented in this work, ranging from SRA samples to several collections of assembled and reference genomes. In addition to user interaction with the web interface, MetaGraph Online provides a web API that allows connecting to the respective servers via their endpoints. That is, any of the hosted indexes can be queried via Python API by connecting to the respective endpoint of the server hosting that index (**Supplemental Figure S-8**). Moreover, indexes constructed from public-access data can be downloaded for large-scale local analyses.

### MetaGraph provides a clear cost advantage

The presented MetaGraph approach is based on the strategy of pre-indexing raw data for accelerated search and optimized storage. While hosting all existing SRA data in a commercial compute cloud would amount to a total cost of $3,951,607 USD per year, storing the MetaGraph indexes even in their most performant representation would decrease the cost by 25% (**Supplemental Table S-5**). In an on-demand setting, the scenario with the least cost overhead, the annual hosting cost reduces to $52,261 (a 75-fold reduction). While existing on-demand approaches such as Serratus^62^ have no hosting cost, every queried Mbp of sequence costs *∼*$1,736. MetaGraph on-demand, on the other hand, reduces this query cost to $2.80 per Mbp for alignment and $0.10 per Mbp for exact *k*-mer matching, resulting in a break-even at an annual query size of 30 Mbp. We have determined the additional cost to index each sample at $0.022, which is negligible compared to costs for data acquisition. Thus, MetaGraph is the hitherto most cost-effective solution to realize SRA full-text search. Full details for our calculations are available in **Supplemental Section A.5**.

## Discussion

In this work, we set out to demonstrate the practical feasibility of indexing whole sequencing archives, such as NCBI’s Sequence Read Archive (SRA), with the goal of making them accessible for full-text search. The major challenge of this task is to hold pace with the exponentially growing amount of input data. As a solution, we have presented MetaGraph, a highly scalable and modularized framework designed to index and analyze very large collections of biological sequence data.

In total we have processed almost 5 Petabases of public sequencing data and have transformed them into compressed indexes that are accessible for full-text search, reducing the input data size up to a factor of *∼*7,400 and showing up to 20-fold reduction compared to current state-of-the-art methods. This not only makes data better accessible but also easily transportable across analysis sites. Specifically, we have indexed various collections of real DNA and RNA sequences, including a significant portion of all publicly available whole-genome sequencing samples from the NCBI SRA. In particular, we have indexed over 90% of all Microbe, Fungi, Plant, Human, human gut metagenome, and a substantial part of the Metazoan samples from the SRA, which together alone make up 2.6 Petabases in 1,903,327 read sets. In addition, we indexed a number of other diverse and biologically relevant data sets, from reference genomes to raw metagenomic reads. The MetaGraph indexes we constructed require orders-of-magnitude less storage than the original gzip-compressed inputs: 4742*×* for GTEx, 77–2595*×* for SRA, 27*×* for MetaSUB, as well as 7.6*×*, 1.6*×*, and 1.1*×* for the UHGG, Tara Oceans MAGs, and RefSeq assembled sequences, respectively.

The scope of our index collection is only limited through the exclusion of controlled-access data and certain long-read sequencing technologies with a very high error rate. In general, we found that the effective handling of sequencing errors and technical variation in the input is one of the greatest challenges in building a joint index. For the data we have indexed, we have shown that, on average, more than 90% of the input data can be accurately aligned back to the index, achieving a very high sensitivity to recover the original experiment of a read.

Due to the sheer size of the data and because our indexes perform optimally when present in main memory, we have split up the input into several batches. Although this creates the overhead of running parallel queries to several indexes, this enables us to arbitrarily scale out the index representation. All partitions can be used to emulate a single large index, which is enabled through MetaGraph’s server mode and the accompanying Python API. A second concession we needed to make to accommodate input size is the removal of spurious input *k*-mers. On all data sets (except assembled references), we apply a strategy of moderate cleaning to not accumulate sequencing noise in the final indexes. Although this slightly reduces our sensitivity for query, we generally observe a 10-fold reduction in index size. It is part of our future work to adapt the raw data cleaning and error correction approaches to also be applicable to sequencing platforms with higher error rates, such as PacBio’s SMRT or reads from Oxford Nanopore Technologies.

In addition to reducing the final representation size, a practical full-text index also needs to optimise for query efficiency. For the experiment discovery problem, we have shown that MetaGraph achieves a performance superior to other state-of-the-art methods. Using *k*-mer based query, we can search for *∼*300 Mbp of sequences per hour per thread against a cohort-size index. Through internal parallelization, rates of 10 Gbp/hour on a single server are easily achievable. Naturally, there is a trade-off between the choice of *k*, the distance between query and target, and the resulting query sensitivity. In settings where an increased sensitivity is needed or the distance between the query and the target is large, we provide several sequence-to-graph alignment strategies. Although this comes at the cost of greater querying complexity, our implementation achieves query times that are only a small constant factor worse than the less sensitive exact matching. The main challenge in the alignment regime is thus balancing the trade-off between alignment search time and sensitivity. In general, there is a trade-off between representation size and query efficiency. MetaGraph addresses this by representing input data as collections of *k*-mer sets stored in various succinct data structures, offering practically relevant trade-offs between index size and query performance. This flexibility allows for running MetaGraph at different scales and on different hardware, from laptops to research compute clusters to distributed cloud environments. As new graph and annotation representations are developed, MetaGraph’s extensible framework allows us to readily integrate these new developments. For example, recent graph data structures with improved data locality^63^ can enable parts of the index to be offloaded to disk to allow for streaming access patterns during search. Notably, all existing indexes can be easily transformed into new formats, directly benefiting from algorithmic improvements in the field.

We provide all indexes generated from public data as an open resource to the community. Through providing a web service for interactive queries at MetaGraph Online, we allow life science researchers and other communities, at last, easy and cost-efficient access to the sequence data for exploration and search. Although this still only represents about 10% of all available sequencing data, our effort has been carried out by a single research lab, demonstrating general feasibility and serving as a proof of concept. Processing the remaining 90%, as well as all future incoming data, can be easily solved via scaling through parallelisation and should be quite feasible for institutions such as EBI or NCBI.

The sequence search results we present in this work set a crucial milestone in computational modelling, solving the problem of making all existing biological sequence data searchable and easily accessible for exploration – a very timely problem of high practical relevance. For the first time, it is now possible to efficiently search for nucleotide sequences in close to all sequenced raw genomes across the tree of life. What was deemed challenging a few years ago, such as indexing and searching in a few thousand read sets, now is tractable and can easily be done on a modern laptop. We envision MetaGraph to serve as a versatile tool that enables researchers to perform large-scale comparative analyses in genomics and medicine on typical academic compute clusters, making public data sets truly open and interactively accessible. In particular, MetaGraph indexes can facilitate large-scale learning tasks on biological sequences (e.g., DNA, RNA, amino acid), such as training large language models.

The size and the diversity of the data we have processed prove the feasibility of keeping all existing sequence archives indexed in a general manner and making them searchable, similar to how Google indexes web pages and the information extracted from them. We emphasize that the support for sequence alignment is especially crucial, for instance, when searching for highly dissimilar sequences with high mutation rates. Together with its remarkable scalability, this makes MetaGraph stand out from all state-of-the-art methods and tools.

In addition to the basic search functionality, the MetaGraph framework allows for sequence assembly from sub-graphs, currently enabling our graph cleaning approach. A goal for improving this functionality is increasing contig lengths. While many sequence assemblers take an iterative cleaning approach to remove noise and produce longer contigs^33,64,65^, the alternative approach of chaining together unitigs to form longer contigs is a clear target for future work^66^. In particular, the use of the annotation matrix to guide the path traversal strategy could increase contig lengths without compromising assembly quality with heavy cleaning procedures.

An exciting new possibility enabled by our assembly framework is the ability to fetch biological sequences specific to certain properties or groups of interest (e.g., individual samples, patient subgroups, or any set function of them). This generalized view on assembly is an extension of pure selection of differential *k*-mers^67^ and allows for new kinds of integrative analyses. For example, MetaGraph can answer such queries as “get all sequences found in samples *x* and *y* but not present in sample *z*”. Theoretically, any logical expression over labels is allowed.

Lastly, MetaGraph has been developed in the context of a highly active research community. Its design as a modular framework allows us to benefit also from future technological improvements. Many new exciting approaches for representing *k*-mer sets^43,63^, the use of approximate-membership-query data structures^34,68–70^, dynamically changing the *k*-mer set^71^, improved annotation compression^72,73^, as well as alternative alignment and seeding approaches^41,74–76^ will be interesting avenues to explore for a future extension of MetaGraph.

We are confident that the approaches presented in this work can be employed and integrated into the infrastructure of large data repositories, such as ENA and NCBI to make all sequence data stored in these repositories searchable, thereby providing essentially a "Google for DNA". Notably, the cost of providing this whole service would be relatively small compared to the price paid to generate this data in the first place and the cost of storing it in the archives.

**Extended Data Figure 1:**
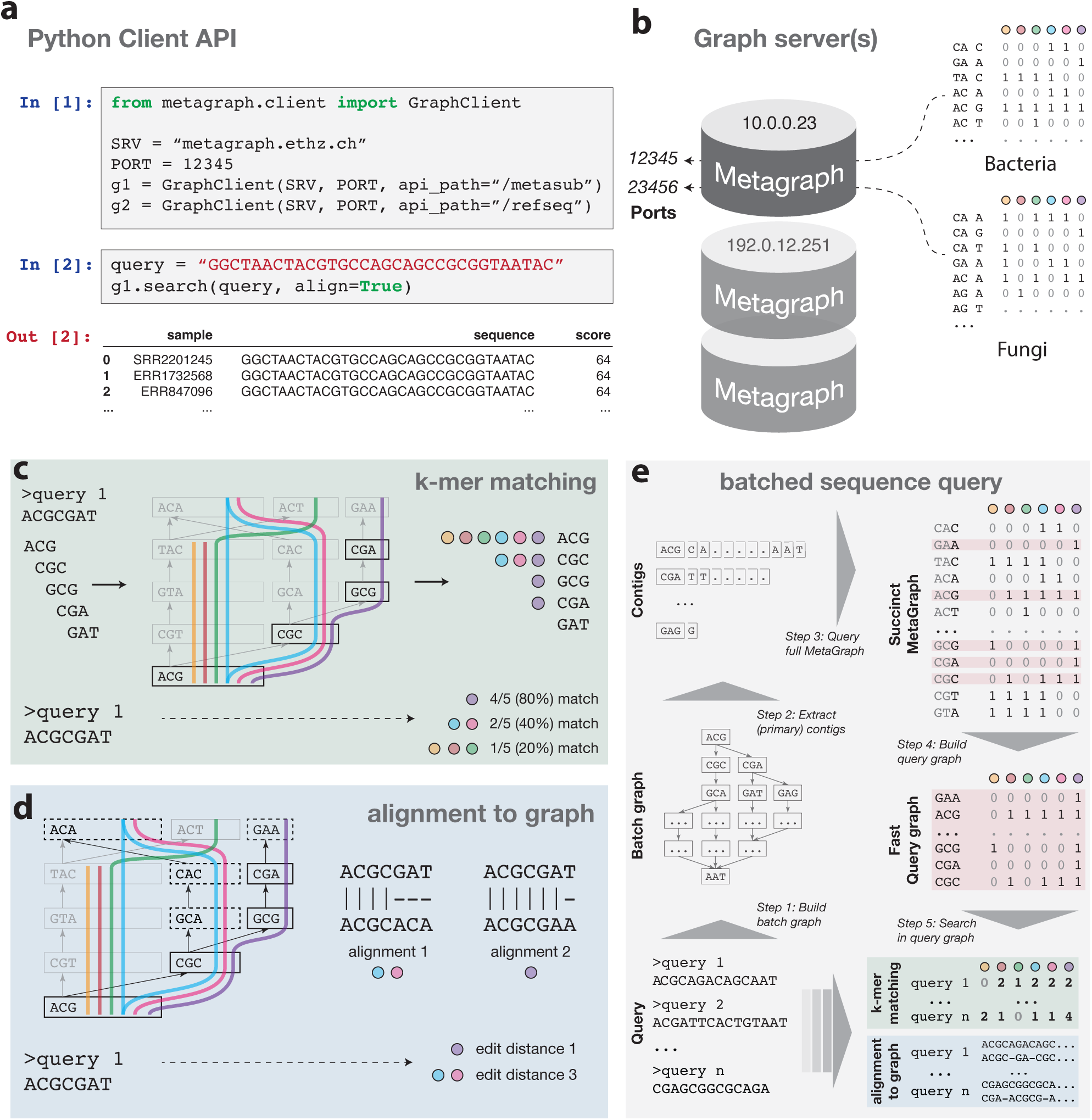
MetaGraph API and search approaches. **a)** MetaGraph is designed to support a client-server infrastructure as exemplified here with a script in Python. In a few steps, several remote (or local) graph instances can be created and queried interactively. Results are returned as a data frame that can be used for further analyses. **b)** Conceptual overview of remote or local index distribution. Every graph index runs on a separate server, accepting queries via the client API. **c-e)** Schematic representation of two main approaches for sequence search. **c)** Counting exact k-mer matches between query and graph. **d)** Alignment finds all closest paths within a given edit distance. **e)** Batched sequence search retrieves a decompressed subgraph (query graph) from the full compressed annotated graph for subsequent query. All query sequences are combined into an intermediate batch graph, which is then traversed to extract contigs to be queried against the full index. Hits and their corresponding annotations are aggregated to construct the final query graph, which is then searched against with the original query sequences.

**Extended Data Figure 2:**
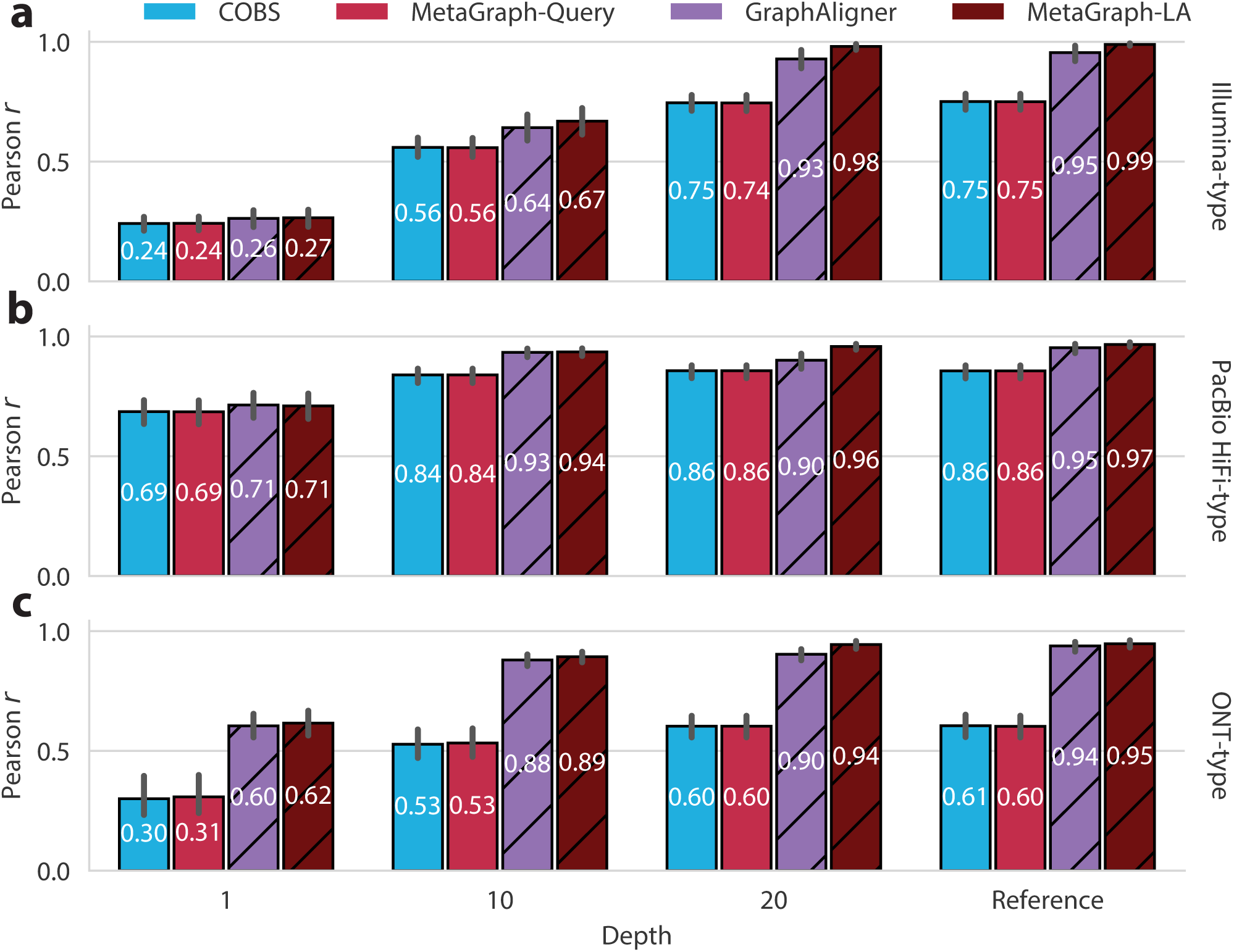
**Accuracy of sequence search approaches** for queries of **a)** Illumina-type, **b)** PacBio HiFi-type, and **c)** ONT-type simulated reads. All graphs (indexes) were constructed from an *Escherichia coli* reference genome or its simulated Illumina-type reads at different sequencing depths. Accuracy is measured as the mean Pearson correlation between the edit distance computed by each method and gold-standard edit distances, measured across 1000 bootstrap samples of 500 simulated *E. coli* query reads. Error bars represent 95% confidence intervals of the mean. Bar hatching indicates a method that uses sequence-to-graph alignment instead of exact k-mer matching.

**Extended Data Figure 3:**
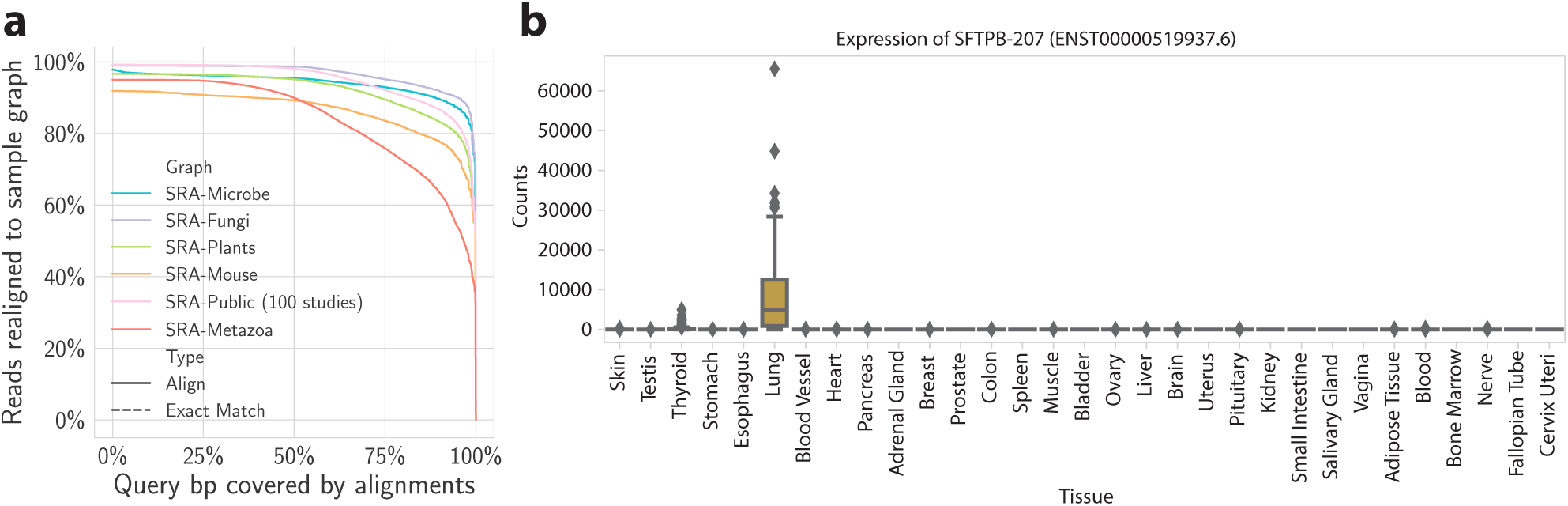
Completeness and specificity. **a)** Complementary ECDF curve showing the fraction of reads that can be re-aligned to the index (y-axis) requiring at least x% matching positions (x-axis) for a range of indexed SRA datasets. **b)** Distribution of sample counts retrieved from the MetaGraph GTEx index for the transcript ENST00000519937.6 of the gene SFTPB, known to be specifically expressed in lung and thyroid tissues.

**Extended Data Figure 4:**
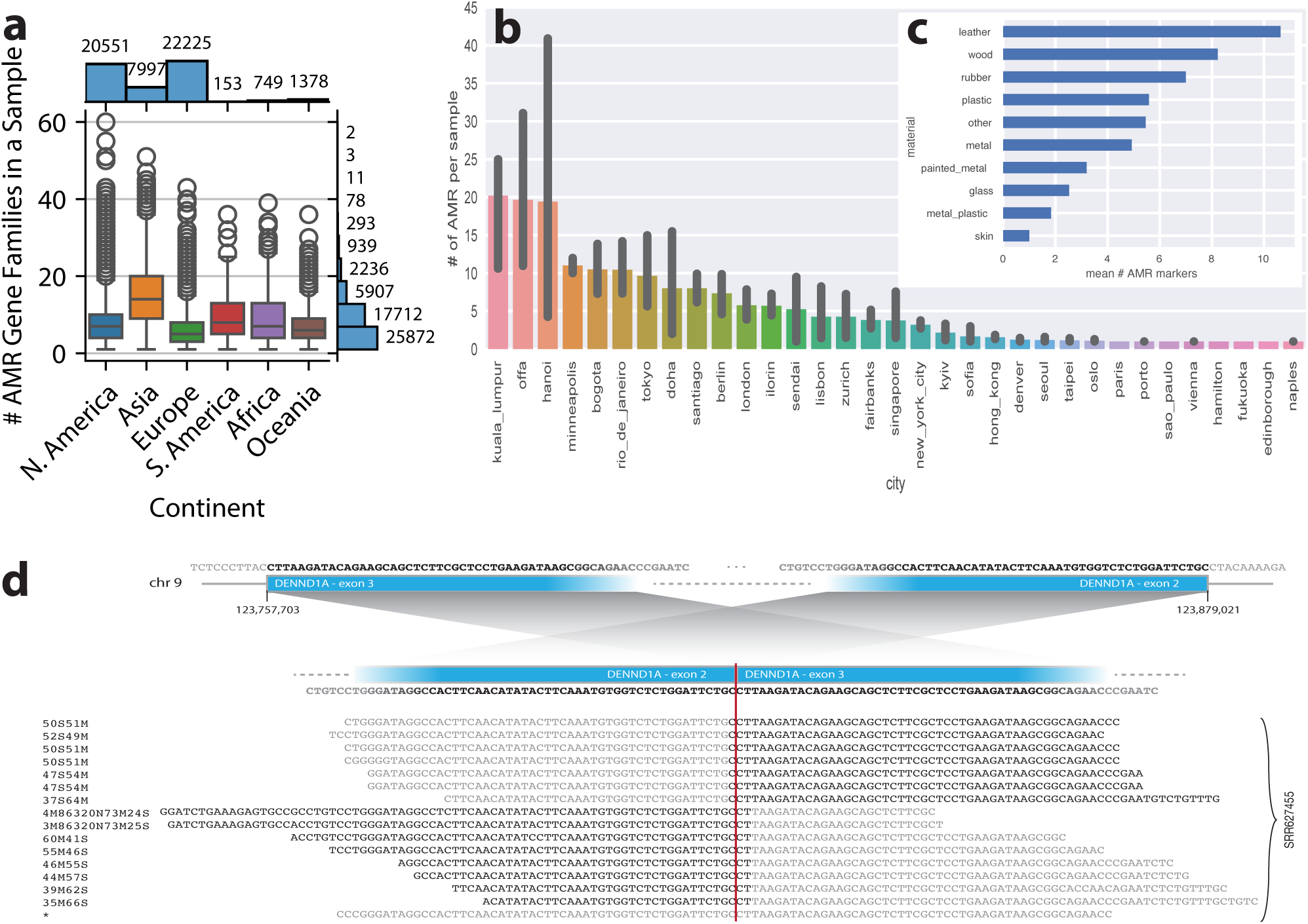
Additional applications. **a)** Number of antimicrobial resistance (AMR) gene families per sample in the SRA-MetaGut human gut microbiome samples. **b)** Number of AMR markers per sample for different cities in the MetaSUB study. Bars represent *±σ*. **c)** Distribution of the mean number of AMR markers grouped by surface material based on all samples of the MetaSUB data set **d)** Back-splice junction detection on the DENND1A gene in GTEx RNA-Seq sample `SRR627455`.

## Methods

### Representing De Bruijn graphs

According to the formal definition of a De Bruijn graph, every two k-mers (short contiguous subsequences of length *k*) with an overlap of *k −* 1 characters are connected with a directed edge. It can be seen that in the De Bruijn graph, the edges can always be derived from the nodes alone, and hence, it is sufficient to store only the k-mer set. These k-mers are used as elementary indexing tokens in MetaGraph.

MetaGraph provides several data structures for storing k-mer sets, which are used as a basis to implement different representations of the *De Bruijn graph* abstraction. In addition to the simple hash table, the k-mers may be stored in an indicator bitmap^77^ (a binary vector represented as a succinct bitmap of size *|*Σ*|^k^* indicating which k-mers are present in the set), or in the BOSS table^3^ (a data structure proposed by **B**owe, **O**nodera, **S**adakane, and **S**hibuya for storing a set of k-mers succinctly). We call our De Bruijn graph implementations based on these data structures `HashDBG, BitmapDBG`, and `Succinct-DBG`, respectively. (For a detailed description of these representations, refer to Supplemental Section A.3.1.) All these data structures support exact membership queries, and they map k-mers to positive indexes from 1 to *n*, where *n* is the number of k-mers in the represented set (or zero if the queried k-mer does not belong to the set). While `HashDBG` is mostly used internally (e.g., for batched sequence search), `SuccinctDBG` typically exhibits the best compression performance, and, thus, serves as the default compressed representation.

### Scalable k-mer enumeration and counting

To count and de-duplicate k-mers, MetaGraph uses the following approach to generate the so-called *k-mer spectrum*: the input k-mers are appended to a list, which is sorted and de-duplicated every time it reaches the allocated space limit or after all input k-mers have been processed. During de-duplication, the k-mer counts are summed up to maintain the total count of each unique k-mer. We call this approach *SortedSet*. In addition, MetaGraph offers the *SortedSetDisk* approach, which implements a similar algorithm in external memory. Using a pre-allocated fixed-size buffer limits memory usage and allows for constructing virtually arbitrarily large k-mer spectra but requires a larger amount of disk I/O. Lastly, MetaGraph supports passing precomputed outputs from the KMC3^78^ k-mer counting tool as input to make use of its exceptionally efficient counting algorithm and filters. Once the entire k-mer spectrum is obtained and all k-mers are sorted, they are converted into the final data structure to construct the target graph representation.

### Extracting contigs and unitigs from graphs

All sequences encoded in the graph (or any defined subgraph) can be extracted from it and stored in FASTA format via graph traversal. The graph is fully traversed and its paths formed by consecutive overlapping k-mers are converted into sequences (*contigs*) that are returned as a result of this operation. Each k-mer of the graph (or subgraph) appears in the assembled contigs exactly once. Thus, the resulting set of sequences is a disjoint node cover of the traversed graph.

MetaGraph provides efficient parallel algorithms for sequence extraction and distinguishes two main types of traversal: i) traversal in *contig* mode extends a traversed path until no further outgoing edge is present or if all the next outgoing edges have already been traversed, while ii) traversal in *unitig* mode only extends a path if its last node has a single outgoing edge, and this edge is the single edge incoming to its target node. This definition of a unitig matches the one from^4^.

### Basic, canonical, and primary graphs

When indexing raw reads sequenced from unknown strands, we supplement each sequence with its reverse complement, which is then indexed along with the original sequence. As a result, the De Bruijn graph accumulates each k-mer in both orientations. Such graphs (which we call *canonical*) can be represented by storing only one orientation of each k-mer and simulating the full canonical graph on-the-fly (e.g., for querying outgoing edges, return not only the edges outgoing from the source k-mer, but also all edges incoming to its reverse complement k-mer).

Storing only *canonical k-mers* (i.e., the lexicographically smallest of the k-mer and its reverse complement) effectively reduces the size of the graph by up to two times. However, this cannot be efficiently used with the succinct graph representation based on the BOSS table. The BOSS table, by design, requires that each k-mer in it has other k-mers overlapping its prefix and suffix of length *k −* 1 (at least one incoming and one outgoing edge in the De Bruijn graph). However, it is often the case that among two consecutive k-mers in a read, only one of them is canonical. Thus, storing only canonical k-mers in the BOSS table would often require adding several extra dummy k-mers for each real k-mer, which makes this approach memory-inefficient. We overcome this issue by constructing so-called *primary graphs*, where the word *primary* reflects the traversal order, as described in the next paragraph.

When traversing a canonical De Bruijn graph, we can additionally apply the constraint that only one of the orientations of a given k-mer is called. More precisely, the traversal algorithm works as usual, but never visits a k-mer if its reverse complement had already been visited. Whichever orientation of the forward or reverse complement k-mer is visited first is considered to be the *primary k-mer* of the pair (for an example illustration see **Supplemental Figure S-5**). This results in a set of sequences, which we call *primary* (*primary contigs* or *primary unitigs*). Note that the traversal order of the graph may change the set of primary sequences extracted from it, but it may never change the total number of k-mers in these sequences (*primary k-mers*). This is relevant when extracting primary contigs with multiple threads since the node traversal order may differ between runs. We call graphs constructed from primary sequences *primary graphs*. Unlike the common approach where only canonical k-mers are stored, primary De Bruijn graphs can be efficiently represented succinctly using the BOSS table, and effectively allow us to reduce the size of the graph part of the MetaGraph index by up to two times.

### Graph cleaning

When a graph is constructed from raw sequencing data, it might contain a considerable number of k-mers resulting from sequencing errors (erroneous k-mers). These k-mers do not occur in the biological sequences and make up spurious paths in the graph, which one may desire to prune off. We call this procedure *graph cleaning*.

MetaGraph provides routines for graph cleaning and k-mer filtering based on the assumption that k-mers with relatively low abundance (low k-mer counts) in the input data were likely generated due to sequencing errors and, hence, should be dropped. To identify potentially erroneous k-mers, we use an algorithm proposed by Iqbal and colleagues^4^. In MetaGraph, we adapted and scaled up this algorithm to work not only for small but also for very large graphs (up to trillions of nodes).

Briefly, to make the cleaning procedure more robust, the decision about filtering out a k-mer is based on the median abundance of the unitig to which this k-mer belongs. That is, k-mers with low abundance are preserved if they are situated in a unitig with sufficiently many (more precisely, at least 50%) highly abundant (*solid*) k-mers. Then the entire unitig is considered *solid* and it is kept in the graph. All solid unitigs (which may also be concatenated into contigs called *clean contigs*) are extracted from the graph and output in FASTA format. Optionally, all tips (i.e., unitigs where the last node has no outgoing edges) that are shorter than a given cutoff (typically 2*k*) are discarded as well. Afterwards, a new graph can be constructed from these clean contigs, which we call a *cleaned graph*.

The abundance threshold for *solid* k-mers (i.e., k-mers that are considered likely non-erroneous) can be set either manually or computed automatically from the full *k-mer spectrum*, where it is assumed that the number of k-mers with sequencing errors (erroneous k-mers) follows a Poisson distribution with a Gamma distributed mean. Also, it is assumed that all k-mers with an abundance of 3 and less are generated due to sequencing errors. Based on these numbers, the algorithm fits a Poisson distribution and chooses a threshold such that k-mers predicted to be erroneous make up at most 0.1% of the total k-mer coverage. Finally, in case the chosen threshold leads to preserving less than 20% of the total coverage, the automatic estimation procedure is deemed unsuccessful and a pre-defined value (typically 2) is used as a fallback threshold instead.

### Constructing a joint graph from multiple samples

According to our workflow, when indexing multiple read sets (especially when indexing vast collections of raw sequence data), the recommended workflow for constructing a joint De Bruijn graph from the input samples consists of the following three steps. First, we independently construct a De Bruijn graph from each input read set. Since each graph is constructed from a single read set (or sample), we call these graphs *sample graphs*. If desired, these sample graphs are independently cleaned with the graph cleaning procedure described above. Then, each sample graph is decomposed into a set of (clean) contigs, either by extracting the contigs directly or as a result of the graph cleaning procedure. Finally, a new De Bruijn graph is constructed from all these contigs, which is then annotated to represent the relation between the k-mers and the input samples. As this graph represents the result of merging all sample graphs, we refer to it as the *joint De Bruijn graph*. In practice, the size of the contigs extracted from sample graphs is up to 100 times smaller than the raw input, which makes the construction of the joint De Bruijn graph by this workflow significantly more efficient compared to constructing it directly from the original raw read sets.

### Graph annotations

Once a De Bruijn graph is constructed, it can already be used to answer k-mer membership queries, that is, to check if a certain k-mer belongs to the graph or not. However, the De Bruijn graph alone can encode no additional metadata (such as sample ID, organism, chromosome number, expression level, or geographical location). Thus, we supplement the De Bruijn graph with another data structure called an *annotation matrix*. Each column of the annotation matrix 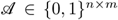, where *n* is the number of k-mers in the graph and *m* is the number of annotation labels, is a bit vector indicating which k-mers possess a particular property:

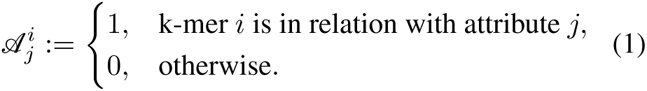

Without loss of generality, we will assume that the annotation matrix encodes the membership of k-mers to different samples, i.e., encodes sample IDs. In this case, 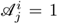 indicates that k-mer *i* appears in sample *j*. (Note that the same k-mer may also appear in multiple samples.)

The number of rows of the annotation matrix corresponds to the number of k-mers indexed in the De Bruijn graph, and hence, the annotation matrix can be of enormous size, containing up to 10^12^ rows and 10^9^ columns. However, this matrix is typically extremely sparse and, thus, can be efficiently compressed.

### Representing graph annotations in MetaGraph

Independent of the choice of graph representation, a variety of methods are provided in MetaGraph for compressing annotation matrices to accommodate different query types. These different matrix representation schemes can be split into row-major and column-major. The row-major representations (`RowSparse`^79^, `RowFlat`^31^, etc.) enable fast row queries but have a poor performance of column queries. The column-major representations (`ColumnCompressed, Multi-BRWT`^11^, etc.), on the contrary, provide fast access to individual columns of the annotation matrix but typically have a poorer row query performance. Despite being mathematically equivalent, up to the transposition of the represented matrix, these schemes are, in fact, different due to the need to adapt to the fact that the number of rows of the annotation matrix in practice is typically orders of magnitude larger than the number of columns. For a detailed description of these and other representation schemes, refer to Supplemental Section A.4.

### The RowDiff compression technique

Due to the nature of De Bruijn graphs and the fact that adjacent nodes (k-mers) usually originate from the same sequences, it turns out that, in practice, adjacent nodes in the graph are likely to carry identical or similar annotations. The RowDiff compression technique^79,80^ exploits this regularity by replacing the annotations at nodes with their relative differences. This allows us to significantly sparsify the annotation matrix and, therefore, considerably improve its compressibility. Notably, the transformed annotation matrix can still be represented with any other available scheme, including those described above. MetaGraph provides a scalable implementation of this technique with efficient construction algorithms, which allow applying it to virtually arbitrarily large annotation matrices. The algorithm essentially consists of two parts. First, for each node with at least one outgoing edge, it picks one of the edges and marks its target node as a *successor*. Second, it replaces the original annotations at nodes with their differences to the annotations at their assigned successor nodes. This delta-like transform is applied to all nodes in the graph except for a small subset of them (called *anchors*), which keep their original annotation unchanged and serve to end every path composed of successors and break the recursion when reconstructing the original annotations (e.g., when querying).

### Counting De Bruijn graphs

Finally, MetaGraph supports generalized graph annotations for representing quantitative information such as k-mer positions and their abundances in input sources, encoded with non-binary matrices^40^.

### Annotation construction

The typical workflow for constructing an annotation matrix for a large input set consists of the following steps. After the joint De Bruijn graph has been constructed from the input sequences, we iterate over the different samples (corresponding to the different annotation labels) in parallel and map all k-mers of each sample to the joint graph, generating a single annotation column. To avoid the mapping of identical k-mers multiple times and to prevent the processing of erroneous k-mers (k-mer with sequencing errors), we use the unitigs extracted from the cleaned sample graphs instead of the raw sequences when annotating the graph. This dramatically reduces the annotation construction time, especially when the joint graph is represented with SuccinctDBG, for which the traversal to an adjacent k-mer is several times faster than a k-mer lookup performed from scratch (see Section A.3.2).

Once an annotation matrix has been constructed (typically in the *ColumnCompressed* representation), it can be transformed to any other representation to achieve the desired trade-offs between the representation size and the performance of the required operations. In particular, for sequence search, the recommended workflow is to apply the RowDiff transform on the annotation matrix and then convert the sparsified columns to the *Multi-BRWT* or *RowSparse* representation, depending on the desired speed vs. memory trade-off.

### Dynamic index augmentation and batch updates

Generally, there are three strategies for extending a fully constructed MetaGraph index (a joint De Bruijn graph and its corresponding annotation matrix). First, the batch of new sequences can be indexed separately and that second MetaGraph index can be hosted on the same or on a different server. Then, these two indexes can be queried simultaneously, as it is done for a distributed MetaGraph index.

Second, the graph can be updated directly if it is represented using a dynamic data structure that supports dynamic updates (e.g., SuccinctDBG (dynamic)). Then, the annotation matrix needs to be updated accordingly. This approach allows making instant changes. However, it does not enable large updates because of the limited performance of dynamic data structures^81^.

Finally, for large updates, the existing index can be reconstructed entirely. For the reconstruction, the index is first decomposed into contig buckets, where each bucket stores contigs extracted from the subgraph induced by the respective annotation column. Then, these buckets are augmented with the new data (either by adding the new sequences directly or by pre-constructing sample graphs from the new sequences and adding extracted from them contigs), and a new MetaGraph index is constructed from these augmented buckets. Notably, this approach uses a non-redundant set of contigs and does not require processing raw data from scratch again. Furthermore, instead of extracting contigs from the old index, it is also possible to use the inputs initially used to construct the old index (e.g., the contigs extracted from sample graphs), which can significantly simplify the process.

### Speeding up k-mer matching

For higher k-mer matching throughput, we implemented several techniques to speed up this procedure. First, when mapping k-mers to a primary graph (defined above), each k-mer may generally have to be searched twice (first that k-mer and then its reverse complement). Nevertheless, if a k-mer has been found, there is no need to search for its reverse complement. In fact, it is guaranteed in that case that the reverse complement k-mer would be missing in the graph. However, if a certain k-mer from the query is missing in the graph but its reverse complement is found, it is likely that for the next k-mer from the query sequence, which is adjacent to the current one, the same applies. Thus, in such cases, we directly start the search for the next k-mer by querying the graph with the reverse complement and checking for the original k-mer only if that reverse complement k-mer is not found in the graph.

When the graph is represented with the BOSS table, indexing *k*-mer ranges in the BOSS table (as described in Supplementary Section A.3.2) greatly speeds up k-mer lookups, especially relevant when querying short sequences or arbitrary sequences against a primary graph.

Another optimization consists of querying the annotation matrix in batches, which improves cache locality and removes possible row duplications. To go further and speed up k-mer mapping as well, we developed the batch query algorithm described below in detail.

### Batched sequence search

To increase the throughput of sequence search for large queries (e.g., sets of sequencing reads or long sequences), we have designed an additional batch query algorithm schematically shown in **Extended Data Figure 1e**. The algorithm exploits possible query set redundancy: the presence of k-mers shared between individual queries. More precisely, query sequences are processed in batches and an intermediate *batch graph* is constructed from each batch. This batch graph is then effectively intersected with the large joint graph from the MetaGraph index. The result of this intersection operation forms a relatively small subgraph of the joint graph, which we call a *query graph* and is represented in a fast-to-query uncompressed format (`HashDBG`). In practice, this intersection is performed as follows. First, the batch graph is traversed (Step 2 in **Extended Data Figure 1e**) to extract a non-redundant set of contigs that are afterwards mapped against the joint graph via exact k-mer matching (Step 3) and the respective annotations are extracted from the compressed index accordingly to construct the query graph with its respective annotations representing the intersection of the batch graph with the full MetaGraph index (Step 4). All query sequences from the current batch are then queried against this query graph (Step 5). Depending on the structure of the query data, this algorithm achieves a 10- to 100-fold speedup.

### Sequence search with alignment

For cases where the sensitivity of sequence search via exact k-mer matching is insufficient, we developed several approaches for aligning sequences to the MetaGraph index, a process known as *sequence-to-graph alignment*. Note that each approach has its target use cases and the choice should be made based on the particular application and the problem setting.

Each alignment algorithm takes a classical seed-and-extend approach^82^. Given an input sequence, the seeds are composed by joining consecutive k-mer matches within the graph’s unitigs (called unitig maximal exact matches, or Uni-MEMs^35^). Although, by default, this restricts the seeds to be at least of length *k*, representing the graph with the BOSS table allows for relaxing this restriction by mapping arbitrarily short sequences to suffixes of the k-mers indexed in the graph (as described in Supplementary Section A.3.2).

Each seed is extended in the graph forwards and backwards to produce a complete local alignment. Similarly to how GraphAligner^5,83^ builds on Myers’ algorithm^84^, our extension algorithm is a generalization of the Smith-Waterman-Gotoh local alignment algorithm with affine gap penalties^85^.

We now describe the extension algorithm in more detail. Given a seed, let *s* = *s*_1_ *· · · s_ℓ_* denote the suffix of the query sequence starting from the first character of the seed. We use a dynamic programming table to represent the scores of the best partial alignments. More precisely, each node *v* has three corresponding integer score vectors *S_v_*, *E_v_*, and *F_v_* of size *ℓ*. *S_v_*[*i*] stores the best alignment score of the prefix *s*_1_ *· · · s_i_* ending at node *v*. *E_v_*[*i*] and *F_v_*[*i*] represent the best alignment scores of *s*_1_ *· · · s_i_* ending with an insertion and deletion at node *v*, respectively.

Let *v_S_* be the first node of the seed *S*. We define an *alignment tree* 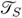 = (*V_S_, E_S_*) rooted at *v_S_* encoding all walks traversed during the search starting from *v_S_*, where *V_S_ ⊂ V ×* ℕ contains all the nodes of the paths originating at *v_S_* and *E_S_ ⊂ V_S_ × V_S_*contains all the edges within these paths. 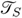 is constructed on-the-fly during the seed extension process by extending it with new nodes and edges after each graph traversal step.

Since the size of 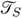 can grow exponentially if all paths are explored (and is, in fact, of infinite size if the graph is cyclic), we traverse the graph and update 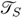 in a score-guided fashion. For this, we maintain a priority queue graph nodes and corresponding score vectors to be traversed, prioritizing nodes whose traversal led to the best local score update^83^. We employ several heuristics to restrict the alignment search space. Firstly, we employ the *X*-drop criterion^86,87^, skipping an element if it is more than *X* units lower than the current best-computed alignment score. Additionally, we maintain an aggregated score column for each graph node storing the element-wise maximum score achieved among the score columns of each node in 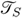. Using this, we discard nodes in 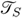 from further consideration if their traversal did not update the aggregate score column. Finally, we apply a restriction on the total number of nodes which can be explored as a constant factor of *ℓ*.

To find seeds of length *< k* matching to the suffixes of nodes in the canonical graph, a three-step approach is taken. First, seeds corresponding to the forward orientation of the query are found according to the algorithm described above. The next two steps then retrieve suffix matches which are represented in their reverse complement form in the graph. In the second step, the reverse complements of the query k-mers are searched to find node ranges corresponding to suffix matches of length *k^′^*. Finally, these ranges are traversed forwards *k − k^′^* steps in the graph to make the prefixes of these nodes correspond to the sequence matched. The reverse complements of these nodes are then returned as the remaining suffix matches.

While primary graphs act as an efficient representation of canonical De Bruijn graphs, special considerations need to be made when aligning to these graphs to ensure that all paths that are present in the corresponding canonical graph are still reachable. For this, we introduce a further extension of the alignment algorithm to allow for alignment to an implicit canonical graph while only keeping a primary graph in memory. During seed extension, the children of a given node are determined simply by finding the children of that node in the primary graph, along with the parents of its reverse complement node. Finding exact matching seeds of length *≥ k* can be achieved in a similar fashion, searching for both the forward and reverse complement of each k-mer in the primary graph.

MetaGraph maintains three different alignment approaches that determine how the graph is traversed during seed extension, called MetaGraph-Align, MetaGraph-LA, and TCG-Aligner, each applying different restrictions to traversal.

#### MetaGraph-Align

In this approach, the sequences are aligned against the joint De Bruijn graph to compute their respective closest walks in the graph. After computing a set of alignments, they are used in place of the original sequences to fetch their corresponding annotations. This approach allows for aligning to paths representing *recombinations* of sequences across annotation labels.

#### Label-consistent graph alignment

When label recombination is not desired, we support an alternative approach where queries are aligned to subgraphs of the joint graph induced by single annotation labels (columns of the annotation matrix). We call this approach *label-consistent* graph alignment (or *alignment to columns*) and is implemented by the MetaGraph-LA algorithm^46^. However, instead of aligning to all the subgraphs independently, we perform the alignment with a single search procedure while keeping track of the annotations corresponding to the alignments.

#### Trace-consistent graph alignment (TCG-Aligner)

Finally, when input sequences are losslessly encoded in a MetaGraph index, the alignment can be done against those original input sequences whose respective walks in the graph are called *traces*. This method is called the *trace-consistent graph aligner* (TCG-Aligner)^40^.

### Column transformations

In addition to the operations mentioned above, MetaGraph supports operations aggregating multiple annotation columns to compute statistics for the k-mers and their counts (abundances). In general, the following formula is used in the aggregation to compute the *i*-th bit of the new annotation column:

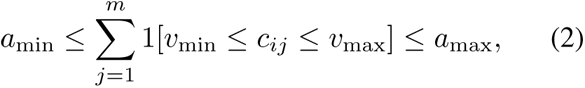

where *c_ij_* is the count (abundance) of the *i*-th k-mer in the *j*-th label, and 1[*A*] is a boolean predicate function, which evaluates as 1 if the statement *A* is true and as 0 otherwise. If no counts are associated with the column, we assume *c_ij_* = 1 for every set bit in the *j*-th annotation column and 0 otherwise. If the sum 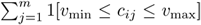 falls within specified minimum and maximum abundance thresholds *a*_min_ and *a*_max_, the bit in the aggregated column for this k-mer is set to 1, and the value of the sum is written as the count associated with that bit. In other words, the resulting aggregated column is always supplemented with a count vector representing the number of original annotation columns with k-mer counts between *v*_min_ and *v*_max_, which can be used in downstream analyses as an ordinary count vector.

### MetaGraph API and Python client

For querying large graph indexes interactively, MetaGraph offers an API that allows clients to send requests to a single or multiple MetaGraph servers. When started in server mode, the MetaGraph index will be persistently present in server memory, which will accept HTTP requests on a pre-defined port. To make the querying more convenient, we have also implemented a Python API client as a Python package available at https://github.com/ratschlab/metagraph/tree/master/metagraph/api/python.

### Indexing public read sets from the NCBI Sequence Read Archive (SRA)

We have split the set of all read sets from SRA (excluding sequencing technologies with high read error rates, see more details in the sections below) into different groups of related samples and constructed a separate MetaGraph index for each group. The groups were defined either using data set definitions of prior work^25^ or using the metadata provided by NCBI SRA. As a result, we constructed the following 6 data sets: SRA-MetaGut, SRA-Microbe, SRA-Fungi, SRA-Plants, SRA-Human, and SRA-Metazoa. All these data sets are listed in **Extended Data Table 1** and make up a total of 4.4 Pbp and 2.3 PB of gzip-compressed input sequences, while the indexes make up only 11.6 TB, which corresponds to the overall compression ratio of 193*×*, or 376 bp/byte.

For constructing the SRA-Microbe index, we used cleaned contigs downloaded from the European Bioinformatics Institute FTP file server provided as Supplementary Data to BIGSI^25^. Thus, no additional data preprocessing was needed for this data set.

For all other data sets, each sample was either transferred and decompressed from NCBI’s mirror on the Google Cloud Platform or, if not available on Google Cloud, downloaded from the ENA onto one of our cloud-compute servers and subjected to k-mer counting with KMC3^78^ to generate the full k-mer spectrum. If the median k-mer count on the spectrum was less than 2, the sample was further processed without any cleaning. Otherwise, the sample was subjected to cleaning with the standard graph cleaning procedure implemented in MetaGraph, with pruning tips shorter than 2*k* (for all these data sets *k* was set to 31) and using an automatically computed k-mer abundance threshold for pruning low-coverage unitigs, with a fallback threshold value of 3. This cleaning procedure was applied for SRA-Fungi, SRA-Plants, SRA-Human, and SRA-Metazoa.

For read sets of the SRA-MetaGut data set, the sequencing depth was typically low, and hence, we applied a more lenient cleaning strategy. Namely, we have switched off the singleton filtering on the k-mer spectrum and used a constant cleaning threshold of 2 during graph cleaning to remove all unitigs with a median k-mer abundance of 1.

For each data set, we first constructed a joint canonical graph with *k* = 31 (including for each indexed k-mer its reverse complement) from the cleaned contigs and then transformed it into a primary graph (storing only one form of each k-mer and representing the other implicitly). Finally, using the same cleaned contigs, we annotated the joint primary graph with sample IDs to construct the annotation matrix. Each input sample thereby formed an individual column of the annotation matrix. The annotation matrix was then transformed to the `RowDiff<MULTI-BRWT>` representation for higher compression and faster queries. The graph was, in turn, transformed to the small representation. For exact commands and scripts refer to Supplemental Data.

### SRA subsets composition

Here we provide a detailed description of each of the 6 data sets.

#### SRA-Microbe

This data set was first used to construct the BIGSI index^25^. Consisting of 446,506 microbial genome sequences, this data set once posed the largest indexed set of raw sequencing data. To the moment of performing our experiments, however, it only represented an outdated snapshot of the corresponding part of the SRA. Nevertheless, we decided to keep the same sequence set for this work to enable direct comparison and benchmarking. A complete list of SRA IDs contained in this set is available as file `TableS1_SRA_Microbe.tsv` (with further information available in `TableS10_SRA_Microbe_McCortex_logs.tsv.` and `TableS11_SRA_Microbe_no_logs.tsv`) in Supplemental Data. For details on how the set of genomes was selected, we refer to the original publication^25^.

#### SRA-Fungi

This data set contains all samples from the SRA assigned to the taxonomic ID `4751` (Fungi) specifying the library source `GENOMIC` and excluding samples using platforms `PACBIO_SMRT` or `OXFORD_NANOPORE`. In total, this amounts to 125,585 samples processed for cleaning. Out of these, 114,839 (91.4%) could be successfully cleaned and were used to assemble the final MetaGraph index. All sample metadata was requested from NCBI SRA on 25.09.2020 using the BigQuery tool on the Google Cloud Platform.

#### SRA-Plants

This data set contains all samples from the SRA assigned to the taxonomic ID `33090` (Viridiplantae), specifying the library source `GENOMIC` and excluding samples using platforms `PACBIO_SMRT` or `OXFORD_NANOPORE`. In total, this amounts to 576,226 samples processed for cleaning. Out of these, 531,736 (92.3%) could be successfully cleaned and were used to assemble the final MetaGraph index. All sample metadata was requested from NCBI SRA on 17.08.2020 using the BigQuery tool on the Google Cloud Platform.

#### SRA-Human

This data set contains all samples of assay type `WGS`, `AMPLICON`, `WXS`, `WGA`, `WCS`, `CLONE`, `POOLCLONE`, or `FINISHING` from the SRA assigned to the taxonomic ID `9606` (Homo sapiens) specifying the library source `GENOMIC` and excluding samples using platforms `PACBIO_SMRT` or `OXFORD_NANOPORE`. In total, this amounts to 454,252 samples processed for cleaning. Out of these, 436,502 (96.1%) could be successfully cleaned and were used to assemble the final Meta-Graph index. All sample metadata was requested from NCBI SRA on 12.12.2020 using the BigQuery tool on the Google Cloud Platform.

#### SRA-Metazoa

This data set contains all samples from the SRA assigned to the taxonomic ID `33208` (Meta-zoa) specifying the library source `GENOMIC` and excluding samples using platforms `PACBIO_SMRT` or `OXFORD_NANOPORE`. In total, this amounts to 906,401 samples processed for cleaning. Out of these, 805,239 (88.8%) could be successfully cleaned and were used to assemble the final MetaGraph index. All sample metadata was requested from NCBI SRA on 17.09.2020 using the BigQuery tool on the Google Cloud Platform.

#### SRA-MetaGut (human gut microbiome)

This group contains all sequencing samples of the assay type `WGS` and `AMPLICON` from the SRA assigned to the taxonomic ID `408170` (Human gut metagenome), excluding samples using platforms `PACBIO_SMRT` and `OXFORD_NANOPORE`. In total, this amounts to 242,619 samples, where 177,759 (73.3%) were `AMPLICON` and 64,860 (26.7%) were `WGS` samples. All these samples were successfully cleaned and were used to assemble the final MetaGraph index. All sample metadata was requested from NCBI SRA on 01.10.2020 using the Big-Query tool on the Google Cloud Platform.

The complete lists of all samples (including the list of successfully cleaned ones) for each subset are available in Supplemental Data.

### Indexing Genotype Tissue Expression (GTEx) data

The 9,759 raw RNA-Seq samples of the Genotype-Tissue Expression (GTEx) project have become a *de facto* reference set for the study of human transcriptomics^50^. All available RNA-Seq samples that were part of the version 7 release of GTEx were downloaded via dbGaP to our compute cluster of ETH Zurich. A list of all samples used is available in the Supplemental Data.

Each sample was individually transformed into a graph using *k* = 31 and then cleaned with the standard graph cleaning algorithm implemented in MetaGraph, with trimming tips shorter than 2*k* and using an automatically computed coverage threshold with the fallback value of 2 for removing unitigs with low median k-mer abundance. All resulting cleaned contigs were assembled into a joint canonical De Bruijn graph and then transformed to the final primary graph. Using the typical workflow, the primary joint graph was annotated using the cleaned contigs extracted from each sample, generating one label per sample. All individual annotation columns were finally collected into one matrix and transformed into the `RowDiff<MULTI-BRWT>` representation.

When performing the indexing with k-mer counts (row ‘GTEx with counts’ in **Table 1**), we applied an additional smoothing of k-mer counts within cleaned unitigs to facilitate the compression. We used a smoothing window of size 60. That is, for each k-mer of a cleaned unitig, its count was replaced with the median abundance of 30 k-mers before it in that unitig and 30 after. This smoothing window is significantly smaller than the expected transcript length, however, it was sufficient to considerably reduce the annotation size (from 184 GB when indexing the original counts to 76 GB).

### Indexing the TCGA RNA-Seq cohort

The Cancer Genome Atlas (TCGA) has collected RNA-Seq samples on the same order of magnitude from primary tumors, spanning across more than 30 cancer types, constituting a central resource for cancer research^49^. We downloaded the data from the Genomic Data Commons Portal of the NCI. A list containing all processed samples is available as file `TableS8_TCGA.tsv.gz` in the Supplemental Data. In total, the index contains 11,095 individual records spanning all available TCGA cancer types. We used the same indexing workflow as for GTEx. Similarly to GTEx, we have also constructed MetaGraph indexes with k-mer counts for TCGA (see **Table 1**).

### Indexing environmental metagenome samples (Meta-SUB)

This data set contains 4,220 whole metagenome sequencing samples collected from the built environment through the MetaSUB consortium^48^. The swabs were collected at different locations and from different objects, where we also contributed to data collection by collecting swabs from benches, ticket machines, and various other objects at different tram stops and train stations in Zurich. When sampling, each swab was annotated with additional data including the location of sampling, the type of object from which the swab was collected, the material of that object, the elevation above or below sea level, and the station or line where the sample was collected. The swabs were then sent for further processing to the sequencing team. For more details about DNA extraction and sequencing, refer to the original publication^48^.

The raw data (read sets) can be downloaded using the MetaSUB utils^88^. A list of all sample IDs used in this study is available as file `TableS6_MetaSUB.csv.gz` in the Supplemental Data.

All input samples were directly assembled into canonical De Bruijn graphs (sample graphs) with *k* = 41. All graphs were then cleaned with the standard graph cleaning procedure implemented in MetaGraph, with pruning tips shorter than 2*k* and removing unitigs depending on coverage (automatically computed based on k-mer spectrum). If no threshold could be computed by the algorithm, we used 3 as a fallback value. The cleaned graphs were transformed into primary contigs, which were then used to assemble a joint graph and annotate it. We annotated the graph with sample IDs, which is the most fine-grained annotation we could construct. Thus, each sample was transformed into a single annotation column in the final MetaGraph index. All the annotation columns were finally aggregated into a joint annotation matrix compressed with the `RowDiff<MULTI-BRWT>` representation. The additional metadata, such as the location, the object, and the surface material, were written to a separate table and could easily be retrieved for any sample ID.

### Indexing the Reference Sequence (RefSeq) collection

The NCBI RefSeq database^52^ contains a non-redundant collection of genomic DNA sequences (all assembled reference genome sequences), transcripts, and proteins.

We indexed all 32,881,422 nucleotide sequences from release 97 of the RefSeq collection, a total of 1.7 Tbp, which takes 483 GB when compressed with gzip −9. We constructed three indexes of different annotation granularity: annotating NCBI Taxonomy IDs at the genus level (85,375 binary annotation columns), annotating sequence accessions (32,881,348 binary annotation columns), and finally annotating k-mer coordinates within buckets split by Taxonomy IDs (85,375 annotation columns with tuples of k-mer coordinates). All these three indexes employ the same underlying De Bruijn graph as a k-mer index with *k* = 31, encoding all the k-mers extracted from the sequences of the RefSeq collection. The graph was constructed in *basic* mode (non-canonical, non-primary), as all the sequences of the collection are assemblies and, hence, are of a determined orientation.

The summary of the indexes is shown in **Table 1**. As expected, the compression ratio (the ratio between the compressed input and the index size) depends on the annotation granularity, varying from 1.9*×* for annotations derived from Taxonomy IDs to 1.0*×* for lossless indexing with k-mer coordinates. Our indexes form an alternative to the commonly used BLAST database^19^ for competitive high-throughput search^40^.

### Indexing global ocean microbiome (Tara Oceans) data

This collection contains 34,815 genomes reconstructed from metagenomic data sets from major oceanographical surveys and time-series studies with high coverage of global ocean microbial communities across ocean basins, depth layers, and time^54^. In addition to metagenome-assembled genomes (MAGs) constructed from 1,038 publicly available metagenomes extracted from ocean water samples collected at 215 globally distributed sampling sites, the collection includes a set of single amplified genomes (SAGs) and reference genome sequences of marine bacteria and archaea from other existing databases. For more details on the data composition, refer to the original publication^54^.

We constructed two indexes for this collection (see summary information in **Table 1**). Both indexes employ a De Bruijn graph over *k* = 31 constructed in the *basic* mode, with 62 million k-mers. The first index annotates k-mers to represent their membership to individual filtered scaffolds reconstructed from the metagenomic samples and takes 1.6*×* less space than the gzip-compressed input. The second index encodes the coordinates of the k-mers within individual assembled genomes and, hence, losslessly represents the input sequences. Notably, it still achieves a compression ratio of 1.2*×*. Lastly, in addition to the two indexes described above, we also indexed the raw assembled scaffolded contigs (360 Gbp) in an annotated graph with 318 million annotation labels. Due to the very large number of annotation columns, unlike the annotation matrices in other indexes typically represented in the Multi-BRWT format, for this index, we represent the RowDiff-transformed annotation matrix in the `RowFlat` format for fast row queries.

### Experiments

This section summarizes the experimental setup for the different results presented in this work.

#### Benchmarking BIGSI and COBS

When indexing subsets of the collection of bacterial and viral genomic read sets^25^ in our evaluation experiments, we used BIGSI^25^ with the same parameters as in the original work^25^ (three hash functions with Bloom filters of size 25 *·* 10^6^) and COBS^26^ in two settings: i) four hash functions and the target false-positive rate of 5%; ii) seven hash functions and the target false-positive rate of 1%.

#### Experiment discovery on SRA graphs

We evaluated each graph using 300 randomly selected samples from their respective input sample sets. To generate a query file for a graph, we randomly selected 100 reads (or the entire read set if fewer than 100 reads are available) from each of the 300 selected samples, resulting in query files of 30’000 reads per graph. To generate auxiliary reads with errors, we selected subsets of the original query sets such that 10 random reads would be selected from each sample. Then, we introduced substitution errors to these reads with probabilities 0.1%, 1%, 2%, 5%, and 10% and insertions-deletions at 10% of the substitution probability using Mutation-Simulator^89^. Given these read sets, we then discarded all reads whose mean *k*-mer multiplicities were below the unitig cleaning thresholds determined by the cleaning procedure (**Methods Subsection “Graph cleaning”**).

We evaluated experiment discovery at two different levels of granularity: *(i)* mapping to individual sample graphs (**Extended Data Figure 3 a**) and *(ii)* mapping to joint annotated graphs (**Figure 3 a,b**). When mapping to joint graphs, we only considered mapping results that retrieved the ground-truth label of each query read. For all granularities, we mapped the reads via both exact *k*-mer matching and label-consistent sequence-to-graph alignment using MetaGraph-LA^46^. We measure how well the reads aligned as the percentage of characters in the query that are covered by at least one reported mapping.

#### Human gut resistome and phageome exploration

We queried all antimicrobial resistance (AMR) genes from the Comprehensive Antibiotic Resistance Database (CARD) database (version 3.2.7)^47,56^ and all bacteriophages from RefSeq Release 218^57^. We selected bacteriophages by selecting all viral sequences with the term `phage` in their header. We mapped these sequences to all accessions in the SRA-MetaGut index representing whole-metagenome sequencing samples. We recorded an accession as a match to a query if at least 80% of the query’s k-mers exactly matched a k-mer in the accession. We then reduced the pools of AMR genes, SRA accessions, and RefSeq bacteriophages to those for which at least one match was found.

To measure the degree of association for each AMR gene family-bacteriophage pair, we computed two binary vectors where each index represents a gut microbiome sample. The first vector indicates the presence of at least one gene match from the gene family and the second vector indicates a match to the phage. We then measure the association using the Matthews correlation coefficient (denoted by Corr_MCC_) if both vectors indicate at least 5 present matches (value `1`) and at least 5 absent matches (value `0`).

When measuring the growth of resistance to antibiotics over time in each continent, we normalized the counts in the confusion matrix before computing Corr_MCC_ to correct for differing numbers of samples deposited in each year and from each continent. If we denote the number of accessions from a continent *C* by *n_C_* and the number of accessions from a year *Y* by *n_Y_*, we normalised the counts by letting each accession from continent *C* and year *Y* contribute a count of 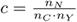 instead of 1. *n_N_* is a scaling factor applied to the four counts in the confusion matrix so that their sum equals the total number of accessions considered. We use the same scaling factor *c* for each count when fitting a linear regression model to the match counts to determine drug resistance growth over time for each continent.

We compute *p*-values for each gene-to-phage correlation Corr_MCC_ and each drug resistance growth linear regression slope *s* via permutation testing. For each analysis, we compute 100 permutations of the antibiotic and gene family indicator vectors, respectively, and compute *p*-values relative to the resulting null distributions.

In **Figure 4 a**, we only plot a gene family or a phage if it has at least one significant correlation with Corr_MCC_ *>* 0.25. In **Figure 4 b**, we report all antibiotics for which we measure statistically significant growth in at least one continent. All *p*-values are corrected using the Benjamini-Yekutieli procedure to a family-wise error rate of 0.05 and are considered to be significant if they are *p <* 0.05 after correction.

#### Re-alignment experiments against GTEx index

We randomly selected a subset of 10 samples from the GTEx cohort (available as file `TableS14_GTExSubset.tsv` in supplemental resources given in Section 1) to evaluate the re-alignment of samples against the graph. For each sample, we considered the first 250,000 read pairs. The reads were re-aligned to the human reference genome (version hg38.p12) using the STAR aligner^90^ (version 2.7.0f). Similarly, we used the MetaGraph sequence-to-graph alignment to align the reads back to the GTEx index. In either setting, we used sensitive alignment criteria, utilizing `-outFilterMatchNmin 21` for STAR and `-align-min-seed-length 21` for MetaGraph. The latter setting does not apply for Meta-Graph exact k-mer matching, where always a full k-mer is mapped.

#### Query of TCGA index

To generate the list of candidate trans-junctions, we iterated over all genes present in the GENCODE annotation (version 32) and formed for all transcripts

### The architecture and the implementation of Meta-Graph Online

The architecture of the MetaGraph Online service is schematically shown in **Figure S-6**. The backend server (**Figure S-6, middle**) implemented as a Flask application provides a web interface and generates dynamic web pages for submitting search queries and viewing the query results. The user queries are processed and transformed into a search query, which is then passed to the remote servers hosting the MetaGraph indexes (**Figure S-6, right**). Once the query has been executed by the respective MetaGraph server, the results are sent back to the backend server and an aggregated summary is rendered to the user. Each MetaGraph index is hosted on a remote server by running the MetaGraph tool in server mode (**Figure S-6, right**). The server applications run independently and are distributed across the available machines. Each MetaGraph server receives HTTP requests formed by the central backend server on user search requests. The communication between the central backend server and the other remote MetaGraph servers happens via the Python API. For seamless compatibility, we also made the backend server redirect user requests and provide the same web API for querying MetaGraph servers (e.g., from Python or as a simple HTTP request) as if the MetaGraph tool is locally running in server mode.

The backend of MetaGraph Online is implemented as a Flask application. This web application is deployed in a Docker container using the Nginx server as a backend. For each search query from the user, it forms a request accordingly and sends it via the Python API client to the MetaGraph servers hosting the indexes. We run these MetaGraph servers in Docker containers on the same or other machines in a federated manner. In addition, the web application emulates the usual MetaGraph API by

## 1 Supplemental Data

Additional resources for this project, including sample metadata, interactive notebooks and analysis scripts are available in GitHub at https://github.com/ratschlab/metagraph_paper_resources. The source code of the MetaGraph software is available under GPLv3 License at https://github.com/ratschlab/metagraph. Besides the core library, the project includes benchmarks, integration_tests for testing the main MetaGraph executable and the command line interface, a set of Snakemake workflows for simplified index construction, a Python API client, and detailed documentation.

## Acknowledgements

This work was supported through funds of ETH Zurich, MK, and HM are funded by the Swiss National Science Foundation Grant No. 407540_167331 “Scalable Genome Graph Data Structures for Metagenomics and Genome Annotation” as part of Swiss National Research Programme (NRP) 75 “Big Data.” The authors would like to thank the members of the MetaSUB international consortium for early access to raw sequencing data and their feedback on early versions of the geolocation DNA sequence search. We also would like to acknowledge the NCBI staff working on maintaining and developing the SRA for their support and interest in our project. The authors thank the members of the Biomedical Informatics Group at ETH Zurich for constant feedback and input on the project. The authors also would like to thank Google Inc. for providing a package of free compute credits on the Google Cloud Platform. We further would like to thank the donors and their families who contributed data to the Genotype-Tissue Expression (GTEx) and The Cancer Genome Atlas (TCGA) projects. This study contains data gathered by the GTEx project available through dbGaP at http://www.ncbi.nlm.nih.gov/gap through dbGaP accession number `phs000424.v7.p1`. This study further contains data gathered by the TCGA project available through dbGaP accession number `phs000178.v1.p1`. The authors are also grateful for the ability to use the publicly available data gathered by the gnomAD project.

## Footnotes

a While Mantis, Bifrost, and MetaGraph provide *lossless* representations of the indexed k-mer sets, BIGSI and COBS both employ probabilistic data structures that lead to false-positive matches.

b `haib18CEM5453_HMCMJCCXY_SL336225`

c Available at https://github.com/ratschlab/metagraph_paper_resources/tree/master/notebooks

