## Supplemental Material for "Indexing All Life’s Known Biological Sequences"

### Chapter A

#### Supplementary Methods

##### A.1 Notation

Let $\Sigma$ be an alphabet of fixed size (e.g., $\{A, C, G, T, N\}$ for DNA sequences and $\{A, R, N, \dots, V, X\}$ for amino acid sequences). Given a choice of alphabet $\Sigma$, we define the extended alphabet $\hat{\Sigma} = \Sigma \cup \{\$$ extended by the sentinel character $\$,$ which is lexicographically smaller than every character of $\Sigma$.

Given a string $s$, let $|s|$ denote its length and let $\varepsilon$ denote the empty string (i.e., $|\varepsilon| = 0$). Let $s_i$ denote the $i$-th character of $s$ and $s_{i:j}$ denote the substring $s_i \cdots s_j$, where $1 \leq i \leq j \leq |s|$. We additionally use the notation $s_{:j}$ to denote the prefix $s_{1:j}$ and $s_{i:}$ to denote the suffix $s_{i:|s|}$. Given another string $s'$ of length $m'$, we denote its concatenation with $s$ by $ss' = s_1 \cdots s_{|s|} s'_1 \cdots s'_{|s'|}$. The power function $c^k$ of a character (string) $c$ defines the concatenation of $k$ characters (strings) $c$. Lexicographical order on strings is denoted by $<$, and co-lexicographical order (i.e., lexicographical order on the reversed strings) by $<^{\text{colex}}$.

Given a positive number $k$, every string of length $k$ is called a $k$-mer. Every $k$-mer over the initial alphabet $\Sigma$ is called a *real $k$-mer* and every $k$-mer containing at least one sentinel character $\$$ from the extended alphabet $\hat{\Sigma}$ is called a *dummy $k$-mer*.

In addition, we use functions $\text{rank}(\cdot, \cdot)$ and $\text{select}(\cdot, \cdot)$ defined for strings in arbitrary alphabets as follows. For a given string $s$, a character $c$, and a positive index $i \geq 1$, function $\text{rank}_c(s, i)$ returns the number of characters $c$ occurring in the string $s$ before index $i$,

$$\text{rank}_c(s, i) = \#\{j \leq i \mid s_j = c\}, \quad (\text{A.1})$$

and function $\text{select}_c(s, i)$ returns the index of the $i$-th occurrence of character $c$ in $s$,

$$\text{select}_c(s, i) = \min\{j \mid \text{rank}_c(s, j) = i\} \quad (\text{A.2})$$

if $i \leq \text{rank}_c(s, |s|)$, and undefined otherwise. For arbitrary arrays and vectors, the functions $\text{rank}(\cdot, \cdot)$ and $\text{select}(\cdot, \cdot)$ are defined similarly as for strings.

##### A.2 Basic building blocks: compressed bitmap representations

In order to minimize the space taken by the MetaGraph indexes while still supporting fast queries, we make heavy use of *compressed data structures*. In this section, we will describe different basic data structures used in MetaGraph for representing bit vectors, which are instrumental in building more complex data structures, such as those encoding graphs and matrices in the MetaGraph index.

##### A.2.1 Schemes for the compressed representation of bit vectors

**Static bit vector.** The fastest of the representations considered here packs the bits in an array of 8-byte integers (stored in type `uint64_t`). In this representation, which we refer to as `stat`, the data is stored uncompressed and therefore takes $n + O(1)$ bits of space, but each bit can be queried in $O(1)$ time, and hence, very quickly in practice. In particular, we used the class `sdsl::bit_vector` from the *sdsl-lite* library [6]. To enable *rank* and *select* operations, two additional data structures have to be added. For these, we used `sdsl::rank_support_v5` and `sdsl::select_support_mcl` data structures from the *sdsl-lite* library, respectively.

`sdsl::rank_support_v5` splits the array into 2048-bit blocks (called *superblocks*) and stores precomputed *rank* values for the last position of each block. This requires 64/2048 bits of space per each represented bit. On top of that, it subdivides each superblock into 5 blocks where the relative *rank* values for each of them are fitted into another 64-bit word as $5 \log_2 2048 = 55 < 64$. Thus, this additional structure requires an extra $128/2048 = 6.25\%$ of additional space while reducing the time complexity of $\text{rank}_1$ to $O(1)$, and more precisely, a few word accesses and popcounts.

`sdsl::select_support_mcl` stores positions of every 4096<sup>th</sup> set bit in an auxiliary vector. These bits are called *sampled bits*. If the distance between a pair of consecutive sampled bits is greater than $\log_2^4 n$, each of the 4096 positions of set bits between them are stored in a packed array using $\log_2 n$ bits per position. This takes at most $\frac{4096 \cdot \log_2 n}{\log_2^4 n} = 4096 / \log_2^3 n$ bits per entry. If the distance between a pair of consecutive sampled bits is less than $\log_2^4 n$, it explicitly stores only the relative positions of every 64-th set bit using less than $\log_2 \log_2^4 n = 4 \log_2 \log_2 n$ bits. This takes, in total, less than $\frac{4 \log_2 \log_2 n}{64}$ bits per bit.

Note that the auxiliary `sdsl::rank_support_v5` and `sdsl::select_support_mcl` data structures are optional and are initialized only when the respective *rank* or *select* operations have to be supported.

**Compressed bit vector *sdarray*.** The *sdarray* compressed data structure was proposed in [11] for representing very sparse bit vectors. It stores the positions of set bits using the Elias-Fano encoding for non-decreasing sequences. The $w := \max(1, \lceil \log_2 n \rceil - \lceil \log_2 m \rceil)$ least significant bits of each position are stored in a packed integer vector, where $n$ is the size of the bit vector and $m$ is the number of set bits in it. Hence, the least significant bits take $mw + O(1)$ bits of space. The remaining most significant bits are represented with a delta coding taking $m + \lfloor n/2^w \rfloor + O(1)$ bits. After simplifications, the total representation size is $m(2 + \log_2 \frac{n}{m})(1 + o(1))$ bits, plus the overhead from two `sdsl::select_support_mcl` data structures enabling the $\text{select}_1$ operation in $O(1)$ time, as well as the $\text{rank}_1$ and the access operations in $O(\log \frac{n}{m})$ time. In practice, we used the `sdsl::sd_vector` class from the *sdsl-lite* library [6]. Since this representation was specifically designed for sparse bit vectors and generates an unreasonably large overhead when representing dense vectors, we flip the bits of the represented bitmap when its density is greater than 0.5.

**RRR succinct bit vector.** Next, we used RRR vectors, which were first proposed in [13] as a succinct data structure for representing bit vectors with constant time query times. Later, several practical improvements were proposed, and finally [10] presented a method reducing the memory overhead and providing excellent time performance for *rank* and *select* queries in practice. We used the implementation of the RRR scheme from the *sdsl-lite* library [6]. One notable parameter of RRR vectors is the block size, which provides a practical trade-off between speed and space overhead. As a result of our preliminary benchmarks, we decided to restrict the choice of available block sizes to two values: 15 and 63, as the improvement of query times for other values comes at too high a cost of the extra space overhead and vice versa.

**Dynamic bit vector.** To support dynamic insertions and deletions, in addition to access, rank, and select queries, we represent bit vectors as B-trees with $B = 16$ children per internal node and each leaf storing a respective 8192-bit block of the represented bitmap [12]. This representation naturally supports insertions and deletions in $O(\log n)$ time.

##### A.2.2 Benchmarks and hybrid bit vector representations

We measured the representation size (**Figure S-1**) and query speed (**Figure S-2**) for each scheme. For this experiment, we generated a series of bit vectors of size $10^9$ and different densities with uniformly distributed set bits. It can be seen that, depending on the density, each scheme has a region where it performs best, either in speed or representation size. Thus, we decided to add three hybrid schemes that would switch between some of the base representation schemes described above, depending on the density of the represented bitmap and the desired query performance and space complexity.

The first scheme is called **smart** and switches between **stat** and **sdarray** (**sd**) representations to provide a very good query performance and, at the same time, take advantage of sparsity and ensure a very good compression for sparse bitmaps.

Next, the **small** representation employs the RRR scheme with a block size of 63 except for the regions of extreme sparsity ($< 5\%$ or $> 95\%$), where **sdarray** achieves a smaller representation size.

Finally, in the **smallrank** scheme, we adaptively switch between **small** and **stat** without the select support data structure. This representation achieves exceptionally good compression while providing a very good query time for access and rank queries.

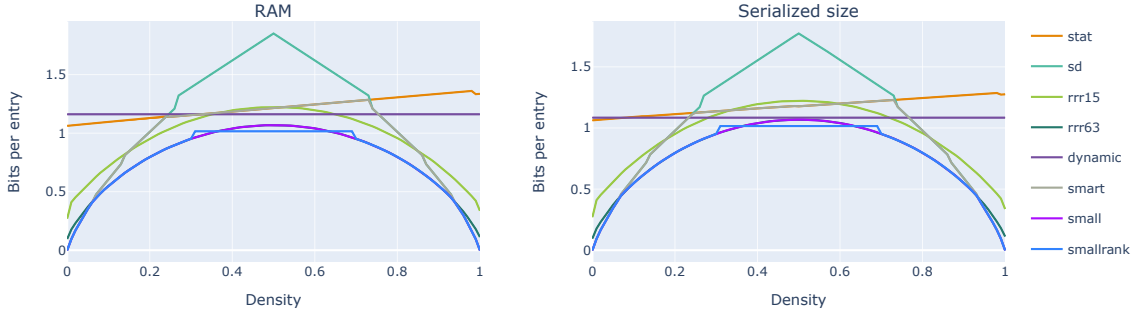

Figure S-1: Size of different bitmap representations (in bits per entry), with set bits uniformly distributed in bitmaps of size $10^9$ and different densities. **Left:** RAM required to load the representation. **Right:** disk space used to store the representation.

#### A.3 Indexing sequences in De Bruijn graphs

By definition, the De Bruijn graph of order $k$ is a pair $(V, E)$, where nodes $V$ are a set of $k$-mers, and directed edges $E$ are all its node pairs $(s_1, s_2) \in V^2$, where the longest nontrivial suffix of the source $k$-mer $s_1$ matches the longest nontrivial prefix of the target $k$-mer $s_2$. The edges $E$ can be unambiguously derived from the nodes $V$. Thus, to represent a De Bruijn graph, it is sufficient to store only its k-mers.

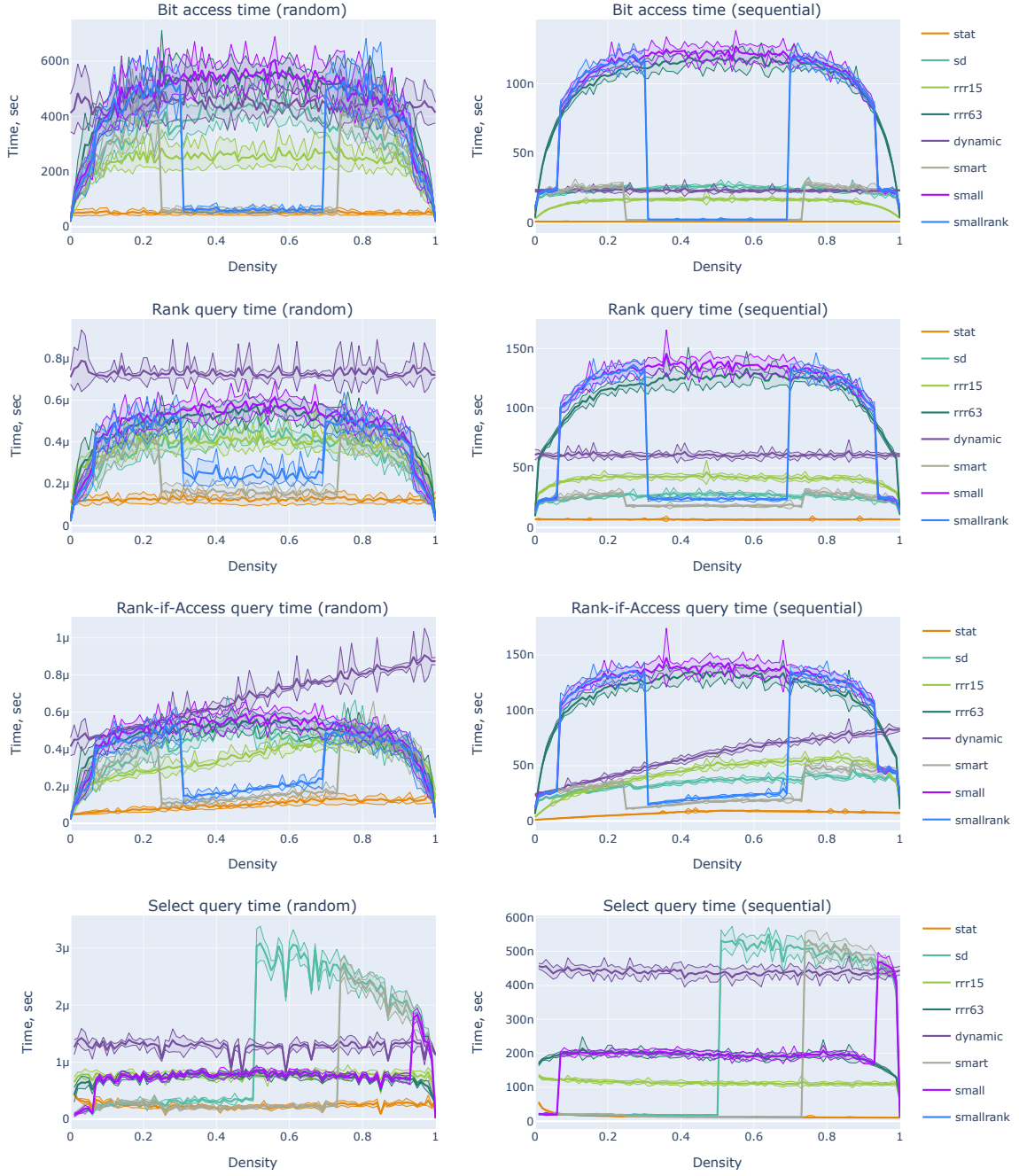

Figure S-2: Average query time for different operations (bit access, rank, rank-if-access, and select queries) for different bitmap representations, with set bits uniformly distributed in bitmaps of size $10^9$ and different densities. **Left:** when queried at random positions. **Right:** average time per query when queried at consecutive positions. The rank-if-access operation combines access and rank and is defined at position $i$ as $\text{rank}(i)$ if $\text{access}(i) = 1$ and zero otherwise.

To index the input sequence data, MetaGraph builds a De Bruijn graph from the input sequences and uses it for two purposes. First, the de Bruijn graph serves as an efficient k-mer dictionary and allows mapping the k-mers onto positive integer indices. Second, the graph topology allows setting the inverse problem of assembling sequences from the k-mers into which the input sequences have been initially decomposed.

##### A.3.1 Representing a De Bruijn graph

MetaGraph employs several data structures (summarized in **Table S-1**) for storing k-mer sets, which are used as a basis to implement different representations of the *De Bruijn graph* abstraction: i) a hash table (**StrHashDBG** and **HashDBG**), ii) an indicator vector (**BitmapDBG**), a binary vector represented as a succinct bitmap of size $|\Sigma|^k$ indicating which k-mers are present in the set [4], and iii) the BOSS table [3] proposed by **Bowe**, **Onodera**, **Sadakane**, and **Shibuya**, storing a set of k-mers succinctly (**SuccinctDBG**). All these data structures support exact membership queries, and they map k-mers to positive indexes from 1 to $n$, where $n$ is the number of k-mers in the represented set (or zero if the queried k-mer does not belong to the set). While **HashDBG** is mostly used internally (e.g., for batched sequence search, see Methods Section “Batched sequence search”), **SuccinctDBG** and **BitmapDBG** exhibit the best compression performance for practical use, depending on the value of $k$. In the next sections below, we describe these data structures in detail.

Table S-1: List of graph representations provided in MetaGraph.

| Graph repr. | $k_{\max}$ | $k_{\max}$ for DNA | Bits per k-mer |
| --- | --- | --- | --- |
| StrHashDBG | unlimited | unlimited | $8k + O(1)$ |
| HashDBG | $\lfloor \frac{256}{\lceil \log_2 \vert \Sigma\vert \rceil} \rfloor$ | 128 | $\max(64, 2^{\lceil \log_2(k \lceil \log_2 \vert \Sigma\vert \rceil) \rceil})$ |
| BitmapDBG | $\lfloor \frac{63}{\lceil \log_2 \vert \Sigma\vert \rceil} \rfloor$ | 31 | $2 + \lceil \log_2 \frac{\vert \Sigma\vert ^k}{n} \rceil + o(1)$ [4] |
| SuccinctDBG | $\lfloor \frac{256}{\lceil \log_2 \vert \Sigma\vert + 1 \rceil} \rfloor$ | 85 | $2 + \log_2 \vert \Sigma\vert + o(1)$ [3] |

###### Hash table-based De Bruijn graph representations

As a basic De Bruijn graph representation, we use a general-purpose hash table to map k-mers packed into 64, 128, or 256-bit integers to their positive integer identifiers (all k-mers are numbered from 1 to $n$). Although this representation is very space-consuming, it enables dynamic insertion and deletion operations and is very useful for algorithm prototyping. In addition, with k-mers stored as strings (with $8k + O(1)$ bits per k-mer), this representation automatically supports k-mers of arbitrary length, which may be important in some applications. As the underlying hash table supports insertions, these representations can naturally be used as dynamic De Bruijn graph representations.

###### Representing a k-mer dictionary with an indicator bitmap

Another De Bruijn graph representation we implemented is **BitmapDBG**, which encodes the presence of k-mers in an indicator vector. Being of size $|\Sigma|^k$ and having set bits in those and only those positions that correspond to the present k-mers, this indicator vector represents a k-mer dictionary and, hence, its respective De Bruijn graph. This scheme was originally proposed in [4]. We store the indicator vector in an **sarray** compressed representation [11], which takes for a vector of size $N$ with $n$ set bits $n(2 + \lceil \log_2 \frac{N}{n} \rceil) + o(n)$ bits of space (or, equivalently, $2 + \lceil \log_2 \frac{|\Sigma|^k}{n} \rceil + o(1)$ bits per k-mer, where $n$ is the total number of k-mers in the dictionary) and performs rank operations in $O(\log \frac{N}{n})$ time, as implemented in the *sdsl-lite* library [6].

##### A.3.2 Succinct k-mer dictionaries represented as a BOSS table

The most scalable and in many ways most versatile De Bruijn graph representation available in MetaGraph is SuccinctDBG. It is based on the succinct self-index proposed by Bowe, Onodera, Sadakane and Shibuya [3] and referred to in the literature as the *BOSS table*. While **Table S-1** only shows the theoretical size of the BOSS table (SuccinctDBG) according to estimations in [3], in practice, we developed three different versions of this representation: SuccinctDBG (static) (for fast queries), SuccinctDBG (dynamic) (supporting k-mer insertions), and SuccinctDBG (small) (a good space vs. time trade-off, in practice typically taking just around 2 or 3 bits per k-mer).

While SuccinctDBG often achieves the best compression of the k-mer dictionary, which becomes a game-changer when indexing data at Petabase scale, this scalability comes at the cost of slower k-mer membership queries to retrieve the respective node indexes in the graph, requiring up to $k$ internal traversal steps [3] for each k-mer query. We alleviate this issue by augmenting the BOSS table with an auxiliary table mapping the k-mer suffixes (usually of length 12) to their respective ranges of rows in the BOSS table (see more details in Section A.3.2). With this optimization and the careful implementation in general, SuccinctDBG achieves a performance, which is sufficient to make the overall query times competitive to other methods while keeping the memory footprint at least an order of magnitude lower (see **Figure 2 a,b**).

In the sections below, we describe this succinct representation in detail, as well as the algorithms for its construction and performing basic operations such as graph traversal and k-mer query.

###### The BOSS representation

The BOSS table [3] consists of three vectors $W$, $F$, and $L$ (defined below) and represents a De Bruijn graph where each node has at least one incoming and at least one outgoing edge (later we show how to loosen this requirement). To satisfy this condition, we extend the alphabet with a special sentinel character $\$$ and add $k$ extra sentinel characters to the beginning and end of every input sequence: $\hat{s} := \$^k s \$^k \in \hat{\Sigma}^* \forall s \in S$. Here $\hat{\Sigma} := \Sigma \cup \{\$\}$ denotes the extended alphabet (i.e., $\hat{\Sigma} = \{\$, A, C, G, T\}$ for DNA) and $\hat{\Sigma}^*$ denotes the set of all finite sequences over this alphabet. As each k-mer, according to this condition, has at least one child and one parent adjacent to it in the De Bruijn graph, we will call a k-mer *first incoming* (*last outgoing*) if there are no colexicographically smaller (larger) k-mers with the same suffix (prefix) of length $k - 1$. (Colexicographic order is obtained by reversing all strings/k-mers, applying lexicographic order, and reversing them again.) The BOSS table requires all k-mers to be sorted in colexicographic order starting from the penultimate character, with ties broken by the last characters. That is, a k-mer $s'_1 \dots s'_k$ precedes another k-mer $s''_1 \dots s''_k$ if k-mer $s'_k s'_1 \dots s'_{k-1}$ is colexicographically less than k-mer $s''_k s''_1 \dots s''_{k-1}$:

$$s'_k s'_1 \dots s'_{k-1} \prec^{\text{colex}} s''_k s''_1 \dots s''_{k-1}, \quad (\text{A.3})$$

which equivalently corresponds to $s'_{k-1} \dots s'_1 s'_k$ being lexicographically smaller than $s''_{k-1} \dots s''_1 s''_k$. Given a sorted list of k-mers $(e^{(1)}, \dots, e^{(n)})$, where $e^{(j)} = e_1^{(j)} \dots e_k^{(j)} \equiv e_{1\dots k-1}^{(j)} e_k^{(j)} \in \hat{\Sigma}^k$, the BOSS

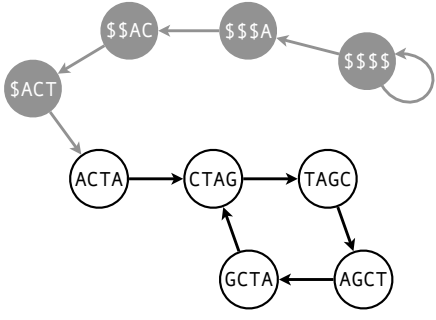

Figure S-3: De Bruijn graph constructed for all k-mers extracted from sequence ACTAGCTAGCTAGC where $k = 4$.

| # | k-mer | $L$ | $F$ | $W$ |
| --- | --- | --- | --- | --- |
| 1 | \$\$\$\$ | 0 | \$ | \$ |
| 2 | \$\$\$A | 1 | \$ | A |
| 3 | \$\$\$AC | 1 | A | C |
| 4 | CTAG | 1 | A | G |
| 5 | \$ACT | 1 | C | T |
| 6 | AGCT | 1 | C | T |
| 7 | TAGC | 1 | G | C |
| 8 | ACTA | 1 | T | A |
| 9 | GCTA | 1 | T | A- |

Table S-2: The BOSS table representation of all k-mers extracted from sequence ACTAGCTAGCTAGC where $k = 4$.

table encodes them in arrays $W'$, $L'$, $L$, and $F$ defined as follows:

$$\begin{aligned} W' &= (e_k^{(1)}, \dots, e_k^{(n)}), \\ L' &= (L'_1, \dots, L'_n), \quad L'_j = \begin{cases} 1 & \text{if } e_{2\dots k}^{(t)} \prec^{\text{colex}} e_{2\dots k}^{(j)} \quad \forall t < j, \\ 0 & \text{otherwise,} \end{cases} \quad j = 1, \dots, n, \\ L &= (L_1, \dots, L_n), \quad L_j = \begin{cases} 1 & \text{if } e_{1\dots k-1}^{(t)} \succ^{\text{colex}} e_{1\dots k-1}^{(j)} \quad \forall t > j, \\ 0 & \text{otherwise,} \end{cases} \quad j = 1, \dots, n, \\ F &= (e_{k-1}^{(1)}, \dots, e_{k-1}^{(n)}). \end{aligned} \tag{A.4}$$

That is, $W' \in \hat{\Sigma}^n$ is the array of the last characters of the indexed k-mers, $L' \in \{0, 1\}^n$ indicates *first incoming* k-mers (see definition above) in the De Bruijn graph, $L \in \{0, 1\}^n$ similarly indicates *last outgoing* k-mers, and $F \in \hat{\Sigma}^n$ is the array of the penultimate characters of the k-mers. Finally, the $W'$ and $L'$ arrays are combined into one vector $W$ as follows:

$$\begin{aligned} W &= ((W'_1, L'_1), \dots, (W'_n, L'_n)) \\ &= ((e_k^{(1)}, L'_1), \dots, (e_k^{(n)}, L'_n)) \in (\hat{\Sigma} \times \{0, 1\})^n, \end{aligned} \tag{A.5}$$

and the vectors $F$, $L$, and $W$ comprise the BOSS representation of the De Bruijn graph corresponding to the list of indexed k-mers $(e^{(1)}, \dots, e^{(n)})$. In practice, however, we extend the alphabet and encode every pair $(c, 0)$, where $c \in \hat{\Sigma}$, by adding a symbol '-' to it (e.g., G-), while every pair $(c, 1)$ is encoded as the same character $c \in \hat{\Sigma}$. As an example, the BOSS table for a De Bruijn graph from **Figure S-3** is shown in **Table S-2**. A demonstration tool for constructing BOSS tables for arbitrary nucleotide sequences is available at https://metagraph.ethz.ch/dbg_visualizer. Note that due to the specific ordering of k-mers in the BOSS table, vector $F$ can be fully represented by $|\Sigma|$ integers, offsets indicating for every $c \in \Sigma$ how many characters lexicographically less than $c$ occur in vector $F$, that is, $F_{\text{offset}}(c) := |\{i \mid F_i < c\}|$.

##### k-mer traversals

In this paragraph, we will show how the BOSS table (i.e., vectors $F$, $L$, and $W$) allows traversing the represented De Bruijn graph along the edges forward and backward.

**Forward traversal.** The problem of *forward traversal* is set as follows. Given an index $j$, $1 \leq j \leq n$, of k-mer $e^{(j)} \in V$ in the BOSS table (where $V \subset \hat{\Sigma}^k$ denotes the set of represented k-mers) and a character $c \in \Sigma$, find index $j'$ associated with its successor (child node) $e' = e_{2,\dots,k}c$ or 0 if such k-mer does not belong to the graph:

$$\text{fwd}(j, c) := \begin{cases} j' : & e^{(j')} = e_{2,\dots,k}c \text{ if } e_{2,\dots,k}^{(j)}c \in V, \\ 0 & \text{otherwise.} \end{cases} \quad (\text{A.6})$$

To execute this operation and find index $j'$, we exploit the specific ordering of the k-mers in the BOSS table. One can always transition to the last outgoing k-mer by computing the index of the last outgoing k-mer for $e^{(j)}$

$$\text{fwd}(j) = \text{select}_1^L \left( \text{rank}_1^L \left( F_{\text{offset}}(e_k^{(j)}) \right) + \text{rank}_{e_k^{(j)}}^W(j) \right), \quad (\text{A.7})$$

and checking the array $W$ for characters $c$ and $c^-$ at positions $\text{select}_1^L(\text{rank}_1^L(\text{fwd}(j) - 1)) + 1, \dots, \text{fwd}(j)$ corresponding to all the k-mers outgoing from $e^{(j)}$ to identify whether one of them ends with character of interest $c$ (see **Table S-2**, e.g., $\text{fwd}(6) = 9$ and $\text{fwd}(6, \mathbf{A}) = 9$). Here $\text{select}_1^L(x)$ returns the position of the $x$-th set bit in array $L$, $\text{rank}_c^W(j)$ returns the number of times character $c$ occurs in array $W$ up to position $j$ (inclusive), and $F_{\text{offset}}(c)$ returns the number of all characters in array $F$ lexicographically smaller than $c$. In practice, we precompute $\text{rank}_1^L(F_{\text{offset}}(c))$ for all $c \in \hat{\Sigma}$ and store these $|\hat{\Sigma}|$ integer values.

**Backward traversal.** We define the *blind backward traversal* as follows. Given a k-mer $e^{(j)} \in V \subset \hat{\Sigma}^k$ and associated with it index $j$ in the BOSS table, find the first incoming k-mer for it and return its associated index

$$\text{bwd}(j) = j', \quad \text{where } e_{2,\dots,k}^{(j')} = e_{1,\dots,k-1}^{(j)} \text{ and } W_{j'} \in \hat{\Sigma}. \quad (\text{A.8})$$

Note that the last condition $W_{j'} \in \hat{\Sigma}$ is equivalent to $L_{j'}^L = 1$ and implies that $j'$ is the first k-mer incoming to k-mer $e^{(j)}$. The existence of the solution follows from the existence of at least one incoming edge for each k-mer of the De Bruijn graph encoded in the BOSS table. One can see that this can be computed on arrays $L$, $F$, and $W$ of the BOSS table as follows:

$$\text{bwd}(j) = \text{select}_{e_{k-1}^{(j)}}^W \left( \text{rank}_1^L(j - 1) + 1 - \text{rank}_1^L \left( F_{\text{offset}}(e_{k-1}^{(j)}) \right) \right). \quad (\text{A.9})$$

Indeed, it is easy to see that $\text{rank}_1^L(\text{fwd}(\text{bwd}(j)) - 1) = \text{rank}_1^L(\text{select}_1^L(\text{rank}_1^L(j - 1) + 1) - 1) = \text{rank}_1^L(j - 1)$, as $\text{rank}_1^A(\text{select}_1^A(\text{rank}_1^A(x) + 1) - 1) = \text{rank}_1^A(x) \forall x, A$. That is, $\text{bwd}(j)$ returns a node, which is mapped by $\text{fwd}(\cdot)$ back to $j$ or to another k-mer that shares with $j$ its source (or, equivalently, has the same prefix of length $k - 1$).

##### Decoding k-mers

Note that given an index $j$ of a k-mer in the BOSS table, one can immediately identify the last character of the k-mer from vector $W$ of the BOSS table (see **Table S-2**). Then, with the blind backward traversal, one can transition to an adjacent incoming k-mer and repeat the same operation to extract the next character of the original k-mer. Repeating this procedure $k$ times reconstructs the original k-mer by decoding all its characters.

##### k-mer lookups

In Section A.3.2, we described how given a node index, one can traverse the De Bruijn graph to transition to one of its adjacent nodes. Here, we will describe the algorithm for k-mer lookup. More precisely, given a k-mer $c_1 \cdots c_k$, find its index in the BOSS table or return 0 if such k-mer is not encoded in the BOSS table. We start with the range of k-mers in the BOSS table that have character $c_1$ in vector $F$. That is, all k-mers with pattern $\cdots * c_1 *$ (recall that vector $F$ in the BOSS table encodes penultimate characters of the k-mers stored in it while vector $W$ encodes the last characters of the k-mers). Thanks to the specific order of the k-mers in the BOSS table, this range is continuous, and thus, we denote it as $[\text{first}_1, \text{last}_1]$ (e.g., $[\text{first}_1, \text{last}_1] = [5, 6]$ for character $C$ in the example from **Table S-2**). Then, we tighten this range by moving pointer $\text{first}_1$ to the next occurrence of $c_2$ in vector $W$, and pointer $\text{last}_1$ to the preceding occurrence of $c_2$ in $W$. Suppose, our range after this step is $[\text{first}'_1, \text{last}'_1] \subseteq [\text{first}_1, \text{last}_1]$. Now we make the forward transition with operation $\text{fwd}$ and transform the range to $[\text{first}_2, \text{last}_2] := [\text{select}_1^L(\text{rank}_1^L(\text{fwd}(\text{first}'_1) - 1)) + 1, \text{fwd}(\text{last}'_1)]$, where $\text{fwd}$ is defined by formula (A.7). As a result, the new range $[\text{first}_2, \text{last}_2]$ corresponds to all the k-mers of pattern $\cdots * c_1 c_2 *$ encoded in the BOSS table. After repeating this procedure $k-3$ more times, we get a range corresponding to k-mers $c_1 c_2 \cdots c_{k-1} *$ and complete the operation by checking $W$ for $c_k$ and $c_k$- within that range to locate the k-mer of interest $c_1 c_2 \cdots c_k$. If during the execution of the algorithm a range $[\text{first}'_i, \text{last}'_i]$ becomes invalid (i.e., $\text{first}'_i > \text{last}'_i$), this implies that the k-mer being searched does not belong to the graph, and hence, we return 0.

Also note that by interrupting this algorithm early, after $k' < k$ iterations, we effectively locate all the k-mers with a given suffix (more precisely, the k-mers of pattern $\cdots * c_1 \cdots c_{k'} *$). We call this operation the *sub-k-mer matching*. Sub-k-mer matching is essential for inexact sequence search algorithms described in the sections below. It could also be efficiently implemented with the BitmapDBG representation of De Bruijn graphs, but it would be impossible to perform with hash table-based representations without an exhaustive search.

##### Indexing k-mer ranges by suffix

At the $t$-th iteration of the k-mer lookup algorithm described above, the current range of nodes corresponds to the k-mers in the BOSS table of pattern $\cdots * c_1 \cdots c_t *$. Thus, if all these ranges were precomputed for all k-mer suffixes $c_1 \cdots c_t \in \Sigma^t$, one could immediately get the respective range for any given k-mer suffix and thereby skip the first $t$ iterations of the lookup algorithm. At worst, this would reduce the query time by a factor of $\frac{k}{k-t}$. In the best case, however, this single lookup in the vector of precomputed k-mer ranges would be enough to find that there are no k-mers in the BOSS table with the given suffix. Hence, it would reduce the entire k-mer lookup algorithm to this single lookup in the vector of precomputed k-mer ranges.

We implemented this idea in MetaGraph and found it to be very effective for speeding up queries on the BOSS table. We call this vector of precomputed ranges an *index of suffix ranges* and represent it with the compressed `sarray` [11] bitmap. More precisely, in this representation, we encode a sparse bitmap $SR \in \{0, 1\}^{1+n+2|\Sigma|^t}$, where $n$ is the size of the BOSS table (the number of k-mers), with set bits at positions indicating the borders of the suffix ranges (hence, of size $1 + n + 2|\Sigma|^t$ with $2|\Sigma|^t$ set bits). Namely, the range of k-mers with the $p$-th suffix (out of all $|\Sigma|^t$ possible suffixes) in the BOSS table can be computed as $\left[ \text{select}_1^{SR}(2p+1) - 2p, \text{select}_1^{SR}(2p+2) - 2p - 1 \right)$. This representation requires only about $2|\Sigma|^t \left( 2 + \log_2 \frac{1+n+2|\Sigma|^t}{2|\Sigma|^t} \right)$ bits, hence, approximately $4 + 2 \log_2 \frac{n}{2|\Sigma|^t}$ bits per integer in the table of ranges when $|\Sigma|^t \ll n$. For instance, with $n = 100 \cdot 10^9$, $t = 12$, and $|\Sigma| = 4$, this makes up about $4 + 2 \log_2 \frac{100 \cdot 10^9}{2 \cdot 4^{12}} \approx 27$ bits per range instead of the 128 bits, which we would need if we stored each range with a pair of 64-bit integers. In addition to the space savings, this representation allows querying the ranges in $O(1)$ time (the complexity of the $\text{select}_1(\cdot)$ operation) as well as performing inverse queries (find a suffix of a $k$-mer given its index in the BOSS table) in $O(\log |\Sigma|^t)$ time. Even though the complexity of this operation is asymptotically the same as $t$ backward traversal steps on the BOSS table, in practice, the constant is much lower, which makes the computation of $k$-mer suffixes this way significantly faster than when directly traversing the BOSS table.

##### Dummy $k$-mers in the BOSS table

Since we only make traversal steps from real $k$-mers encoded in the BOSS table and never traverse forward *dummy sink* $k$-mers (those ending with the sentinel character $\$ \notin \Sigma$), the requirement of the existence of at least one outgoing edge in the represented De Bruijn graph has to be applied only to real $k$-mers, that is, those without sentinel characters $\$$. Thus, for each original real $k$-mer $e$ without adjacent outgoing $k$-mers, we only need to add a single dummy $k$-mer $e_{2,\dots,k}\$$ to the BOSS table (see **Figure S-4**).

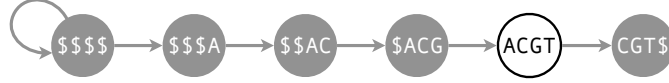

Figure S-4: De Bruijn graph for $k$-mer ACGT with all required non-redundant dummy $k$-mers.

**Redundant dummy $k$-mers and their removal.** Some operations over De Bruijn graphs (e.g., merging two graphs) may lead to the existence of so-called *redundant dummy $k$-mers* in the graph, that is, dummy $k$-mers (with at least one sentinel character) that can be removed from the BOSS table without breaking the properties required for graph traversals. For example, a sink dummy $k$-mer $c_1 \dots c_{k-1}\$$ is redundant if there is another $k$-mer $c_1 \dots c_{k-1}c_k$ that belongs to the graph, where $\$ \neq c_k \in \Sigma$. Redundant dummy source $k$-mers are defined analogously. In MetaGraph, we implemented special procedures to identify and erase these redundant dummy $k$-mers from the BOSS table in linear time.

##### Internal representations of the BOSS table

For different use cases that arise in practice, we developed three different implementations of the BOSS table ($L, F, W$), which we call *graph states*. These different graph states tune the whole graph structure to requirements implied by a particular problem setting and available computational resources. We also present procedures for internal conversion between all the implemented graph states. Given a graph in one state, it can be easily converted to any other.

**Static representation.** In the default static representation, which provides a very good query performance while only taking about $2 + \log_2 |\Sigma|$ bits per $k$-mer, vector $L$ is stored as an uncompressed packed bitmap with the default rank and select support data structures from the *sdsl-lite* library [6] ensuring constant time *rank*, *select*, and *access* operations (see the **stat** bitmap representation in Section A.2), vector $W$ is encoded using the Huffman wavelet tree [8] implemented in the *sdsl-lite* library [6]. Vector $F$, as always, is simply represented as an array of $|\Sigma|$ offsets.

**Small representation.** For the smallest memory footprint, we encode the internal bitmaps of the Huffman wavelet tree $W$ in RRR vectors [13] implemented in the *sdsl-lite* library [6], which ensures

a nearly optimal compression. Similarly, vector $L$ is encoded in a hybrid **small** representation (see Section A.2.2) switching between RRR [13] and **sdarray** [11] vectors, depending on the sparsity of vector $L$ to ensure the best compression. Vector $F$ is encoded by the offset array as described above.

**Dynamic representation.** To support dynamic operations on the BOSS table, we use dynamic bit vector and string representations based on the cache-efficient B-trees implemented in library DYNAMIC [12] to represent vectors $W$ and $L$ with the support for dynamic insertion and deletion operations. Vector $F$ is encoded by the offset array as described above and naturally supports constant-time dynamic operations.

In our workflows, we usually first construct a De Bruijn graph in the SuccinctDBG (static) representation and then convert it to SuccinctDBG (small) once this graph has been annotated and the entire MetaGraph index is constructed and ready to serve queries.

##### A.3.3 Graph construction

While the StrHashDBG and HashDBG graph representations are constructed directly by inserting k-mers into a hash table, the more advanced representations BitmapDBG and SuccinctDBG require a sorted list of k-mers for their construction. Thus, the construction from a set of input sequences proceeds in two steps: 1) k-mer extraction, sorting, and de-duplication; 2) construction of the representation (an indicator bitmap or the BOSS table). In the first stage, each k-mer $e$ extracted from the input sequences, depending on the alphabet size and the k-mer length $k$, is represented as a 64-, 128-, or 256-bit integer, where each character is encoded with exactly $b = \lceil \log_2 |\Sigma| \rceil$ bits ($b = \lceil \log_2 |\hat{\Sigma}| \rceil$ when constructing the BOSS table). The layout of the characters in k-mers is defined accordingly to induce the relative order of the k-mers expected in the target data structure. In the case of the BOSS table, in particular, this is the colexicographic order described in Section A.3.2. Once all k-mers have been de-duplicated and sorted, we construct the final representation, that is, a compressed indicator bitmap for BitmapDBG or vectors $L$, $W$, and $F$ of the BOSS table for SuccinctDBG.

##### Distributed construction

For large data sets where the full set of encoded k-mers may not fit in the amount of RAM available, we employ a distributed construction approach. Each process is assigned a suffix of a fixed length $\ell$ and only k-mers with that suffix are extracted and sorted by that process. The resulting $C_i = (W_i, L_i, F_i)$ tuples from these processes are referred to as *graph chunks*, from which the full graph representation can be constructed through index-wise concatenation. More precisely, given graph chunks $C_1, \dots, C_{|\hat{\Sigma}|^\ell}$ sorted by their respective suffixes, the final graph representation is

$$W := W_1 \cdots W_{|\hat{\Sigma}|^\ell}, \quad L := L_1 \cdots L_{|\hat{\Sigma}|^\ell}, \quad F := \sum_{i=1}^{|\hat{\Sigma}|^\ell} F_i. \quad (\text{A.10})$$

#### A.4 Representing binary matrices in MetaGraph

##### A.4.1 Column-major matrix representations

The default representation of graph annotations in MetaGraph is *ColumnCompressed*, which independently stores columns as compressed bit vectors. More precisely, in the hybrid bit vector representation **smart** (described in Section A.2.2). Being highly compressed for sparse columns, this representation provides easy access to individual columns, which is helpful for filtering (e.g., by sample IDs) and selecting label-induced subgraphs. Furthermore, the *ColumnCompressed* representation provides efficient *access* queries, which makes it an excellent choice for querying individual columns. At the same time, the set bits of a column stored in the **smart** representation can be iterated exceptionally quickly, and thus, the entire matrix can be easily transformed to any other format, including row-major formats, for which the matrix is effectively transposed by blocks.

For higher compression performance and faster row queries, we employ the Multiary Binary Relation Wavelet Tree (*Multi-BRWT*) representation scheme [7] with the index columns stored in the compressed **smallrank** representation (see Section A.2.2), which provides excellent compression while enabling fast rank queries. The *Multi-BRWT* representation scheme typically achieves the best compression in real applications, especially where the columns of the annotation matrix are highly correlated, such as those constructed from sequencing samples corresponding to related organisms.

Generally, when performing row queries on an annotation matrix represented with a column-major scheme, we aggregate the query operations and perform them in batches, which improves the cache locality and significantly improves the query performance.

###### A.4.2 Row-major matrix representations

For fast queries on rows (e.g., for sequence search queries), we provide a number of row-major matrix representations. For the fastest queries of rows, we developed a compressed row-major sparse matrix representation *RowCompressed*, which is employed in query graphs (**Extended Data Figure 1 e**) and additionally supports dynamic operations. It stores the matrix as a vector of vectors, where each row is represented as a `folly::SmallVector<uint32_t>` from the Facebook Open-source Library<sup>1</sup>, storing the column indexes of its set bits, which has the same interface as `std::vector<uint32_t>` but significantly lower memory overhead.

The next compression technique, *RowFlat*, was originally employed in VARI [9]. It concatenates all rows into a single sparse bitmap of size $mn$, where $n$ is the number of rows and $m$ is the number of columns of the annotation matrix, and stores this bitmap in a compressed **sarray** [11] representation.

Then, *RowSparse* writes the column indexes of all set bits into an integer array compressed with the Elias delta coding (implemented in the `sdsl::vlc_vector<>` data structure from the *sdsl-lite* library [6]). Additionally, a bitmap of size $n + d$, where $d$ is the number of set bits in the matrix, is stored in the **small** representation (described in Section A.2.2) to encode the offsets pointing to where each row starts in that integer array.

In *BinRel-WT* [2], a similar approach is used with the difference that the integer array is represented as a wavelet tree, which improves the performance of column queries.

Next, *Rainbowfish* [1] uses the fact that rows of the annotation matrix are often highly duplicated (due to multiple nodes in the graph having the same annotations). To take advantage of that, it builds a dictionary of distinct rows and stores a mapping from original row indexes to their corresponding indexes of distinct rows in the dictionary. Additionally, the more frequent rows are assigned smaller indexes in the dictionary, which makes this mapping more compressible with universal codes. Interestingly, this technique can be generalized and used in combination with any matrix representation scheme. We describe this generalization below.

<sup>1</sup>https://github.com/facebook/folly

##### A.4.3 The Rainbow matrix decomposition technique

It is easy to see that the row de-duplication technique used in *Rainbowfish* [1] can generally be used with any compression scheme used to represent the matrix of distinct rows. Moreover, this matrix does not have to be represented in a row-major order but any column-major representation would be applicable as well. We call this technique the *Rainbow* decomposition, and thus, we call the representations employing it the *Rainbow*-\* representation schemes. For example, the *Rainbow-BRWT* representation de-duplicates the rows with the Rainbow technique and stores the distinct rows in a matrix represented with the compressed *Multi-BRWT* scheme. Note that in this terminology, *Rainbow-RowFlat* refers to the original *Rainbowfish* [1] representation.

#### A.5 Estimating the cost of comprehensive nucleotide search engine

In this section, we will estimate the cost of constructing and hosting MetaGraph indexes, as well as the cost of serving user queries for sequence search. First, we will make general calculations and then make calculations specifically for hosting the whole SRA.

##### A.5.1 General estimation methodology and assumptions for MetaGraph

Suppose we would like to index with MetaGraph a certain large data set (such as the SRA) of size $D$, measured in base pairs. We partition the whole data set into $n$ parts and index them independently. As a result, we get $n$ MetaGraph indexes of size $I_1, \dots, I_n$, which we would host on $n$ machines of a certain type to execute user queries.

Next, we will make the following assumptions.

1. All indexes have the same size $I_1 = \dots = I_n$. Moreover, the total size of the indexes $I := \sum_{i=1}^n I_i$ does not depend on the partitioning of the data set and, in particular, on the number of chunks $n$.
2. The resources required to host an index (memory and local disk space) grow linearly with the index size: $R_M(I) = rI$, $R_L(I) = lI$.
3. The throughput for search and alignment is in inverse dependence with the index size $I$: $S(I) = s/I$, where $S(I)$ denotes either the throughput of k-mer matching or one of the alignment algorithms in MetaGraph running on an index of size $I$ with a single thread.

As an implication of the first assumption, one can index a part of the whole data set of size $D/n$ and derive from the size $I$ of this index the compression ratio $c = \frac{D}{nI}$. Essentially, the assumption says that the compression ratio does not depend on the partitioning of the data set. In effect, MetaGraph does exploit the redundancy across the indexed samples to enhance the compression. Thus, the estimation of $c$ on a relatively small subset (especially randomly selected) generally leads to an underestimated value of $c$. In turn, this leads to overestimating the costs of hosting indexes constructed on larger parts of the data set.

The second assumption comes naturally.

The third assumption typically holds true (see the results in **Figure 2b** as an example). Moreover, if, theoretically, the query time grows superlinearly for a certain type of data, the data set can be partitioned into smaller chunks, which would be indexed independently and queried sequentially to keep the growth of the query time linear.

Suppose each hosting machine has $N$ cores, $M$ RAM, $L$ of local disk space, and costs $C$ per time unit. Then, the minimum number of machines required to host the indexes is $n_{\min} := \lceil \max(\frac{rI}{M}, \frac{U}{L}) \rceil$, where $I = c \cdot D$ is the total size of the indexes (we used the first assumption for this derivation). If all the machines are queried simultaneously, the total throughput of such a system can be up to $s / \frac{I}{n_{\min}} \cdot N = sNn_{\min}/I$. Since the cost of the machines is $n_{\min}C$, the effective query cost is $\frac{n_{\min}C}{sNn_{\min}/I} = \frac{C \cdot I}{sN}$. Note that the effective query cost does not depend on the number of machines $n_{\min}$ used. Indeed, by using more machines, we pay more per time unit, but the task is performed faster. Hence, the total cost is the same.

##### A.5.2 Hosting scenarios

We will consider the following scenarios of hosting preconstructed MetaGraph indexes. On the one hand, the MetaGraph indexes can be constantly loaded in RAM to be ready to serve user queries.

On the other hand, the MetaGraph indexes can be precomputed and stored in a cloud storage with fast access to allow the user to start an arbitrary number of machines on demand and execute the queries on them. Each machine would download a precomputed index from the common storage and would run the query against it. With a sufficiently large query (in practice 100 Mbp is already sufficient), the time of downloading and initializing a MetaGraph index can be neglected as it is far smaller than the actual time taken to perform the query against this index.

The first scenario assumes we reserved a sufficient number of machines to keep all the indexes loaded in their RAM. This ensures that one can execute queries online. However, it implies high costs for keeping those machines running constantly. Alternatively, we could reserve significantly fewer machines and keep the indexes in their local storage. Thus, to serve a query, each machine would sequentially load the indexes from the local storage into RAM and would perform the search against them. Effectively, this is equivalent to the strategy 'on-demand' described above, except that we would keep a fixed number of machines ready to serve user queries at all times. As a result, we could use cheaper instances (for dedicated use instead of on-demand) provided by the cloud provider, as in the first scenario. As a downside, the query throughput would be significantly lower, and the delay before seeing the query results would be significantly higher than in the first scenario, which would nevertheless not affect the effective query costs, assuming the query is performed in sufficiently large batches (as in the on-demand scenario).

##### A.5.3 Cost estimates for sequence search against the entire SRA

In this section, we will estimate the cost of search against the whole of SRA, which, according to the statistics provided by NCBI, comprised in January 2023 almost 40 Pbp or 15.68 PB of sequences with public access. Even simply storing this compressed data costs \$3,951,607 per year<sup>2</sup>, or on average 17 cents per sample per year. The detailed growth statistics for the SRA is shown in **Table S-3**.

To the best of our knowledge, the only other approach used in the past to align sequences against samples from the SRA at a Petabase scale is Serratus [5]. In Serratus, the final costs of search were estimated as \$0.0062 per SRA sample with a 79.8 MB (82.16 Mbp) query [5]. After normalization to estimate the effective cost of search per 1 Mbp of query against the current version of the SRA (23,010,648 samples with public access to the beginning of 2023), we get $\$0.0062 \cdot 23,010,648 / 82.16 = \$1'736$.

Next, we will estimate the cost of indexing the entire SRA with MetaGraph and hosting these indexes to provide a service for sequence search, as well as estimate the effective query costs.

<sup>2</sup>The estimate is based on the rate of \$0.021 per GB for the Amazon S3 Standard storage service (data from https://aws.amazon.com/s3/pricing/, accessed on 10.06.2023).

Table S-3: Growth of the Sequence Read Archive (SRA). For every year, the table shows the amount of data with public access in the SRA at the beginning of that year. The values in parentheses show the amount of data added during that year (the delta). The metadata file with this statistics was collected from NCBI on 12.06.2023.

| Year | # samples | Tbp | Size, TB |
| --- | --- | --- | --- |
| 2008 | 234 (+5,944) | 0.02 (+2.2) | 0.05 (+12.8) |
| 2009 | 6,178 (+16,265) | 2.2 (+9.8) | 12.9 (+15.4) |
| 2010 | 22,443 (+37,637) | 12.1 (+47.0) | 28.2 (+72.9) |
| 2011 | 60,080 (+64,534) | 59.1 (+108.6) | 101.1 (+76.0) |
| 2012 | 124,614 (+132,228) | 167.7 (+218.1) | 177.2 (+133.4) |
| 2013 | 256,842 (+187,457) | 385.8 (+482.8) | 310.6 (+296.0) |
| 2014 | 444,299 (+407,098) | 868.6 (+677.2) | 606.6 (+392.5) |
| 2015 | 851,397 (+666,196) | 1,546 (+1,100) | 999.1 (+659.7) |
| 2016 | 1,517,593 (+1,020,304) | 2,645 (+1,479) | 1,659 (+807.6) |
| 2017 | 2,537,897 (+1,178,709) | 4,125 (+1,923) | 2,466 (+951.6) |
| 2018 | 3,716,606 (+1,880,461) | 6,048 (+2,871) | 3,418 (+1,337) |
| 2019 | 5,597,067 (+2,080,174) | 8,919 (+4,650) | 4,755 (+1,865) |
| 2020 | 7,677,241 (+3,254,730) | 13,569 (+6,435) | 6,620 (+2,547) |
| 2021 | 10,931,971 (+5,684,245) | 20,004 (+8,578) | 9,167 (+2,910) |
| 2022 | 16,616,216 (+6,394,432) | 28,581 (+10,369) | 12,077 (+3,604) |
| 2023 | 23,010,648 | 38,950 | 15,681 |

##### Indexing costs

To estimate the total cost of constructing MetaGraph indexes for the entire SRA, we indexed a set of samples from 100 random studies from the SRA. This set contained 5,184 read sets with a total of 9,579 Gbp (this data set is also shown in **Table 1**). We preprocessed all these samples with the standard cleaning workflow implemented in MetaGraph, also applied when indexing SRA-Fungi, SRA-Plants, SRA-Human, and SRA-Metazoa (see Methods Section “Indexing public read sets from the NCBI Sequence Read Archive (SRA)”). Each cleaning task reserved 4 cores and 40 GB RAM of dedicated compute nodes in the cloud. In total, all these tasks took 1370 cpu-hours. For such tasks, **m1-megamem-96** instances provided by Google Cloud<sup>3</sup> at a rate of \$3.20 per hour for committed use was the cheapest option as of 10.06.2023. Each instance has 192 virtual cores and 1433.6 GiB RAM, and hence, can perform 38 tasks at a time. Thus, the preprocessing of the entire SRA would cost $\$3.20 \cdot 1370/38/9.579 \text{ Tbp} \cdot 38,950 \text{ Tbp} \approx \$469,000$, which corresponds to only 2 cents per sample.

The construction of the final MetaGraph index of these 5,184 cleaned samples on a machine with 32 cores and 75 GB RAM took from 23.6 to 24.8 hours (see **Table S-4** for exact values), depending on the final representation. On a **e2-highmem-16** machine (with 32 virtual cores and 128 GiB RAM) provided by Google Cloud at a rate of \$0.33 per hour for committed use, this computation would cost around $\$0.33 \cdot 24 = \$7.92$. After normalization, we estimate the total indexing cost of preprocessed samples from the entire SRA as $\$7.92/9.579 \text{ Tbp} \cdot 38,950 \text{ Tbp} \approx \$32,000$ or only 14 cents per 100 samples. Thus, the total amortized cost of indexing one raw read set from SRA is 2.2 cents.

##### Estimating characteristics of MetaGraph representations for SRA

We measured key statistics for the MetaGraph indexes constructed in different representations for those 5,184 samples (9,579 Gbp) to estimate the parameters $r$, $l$, $s$ for each representation. (These parameters will be used in the next section to estimate the cost of constructing and hosting the index of the entire SRA.) The results are summarized in **Table S-4**. One can see that the index

<sup>3</sup>https://cloud.google.com/

Table S-4: Key statistics of MetaGraph indexes constructed from the 5,184 samples (the total of 9,579 Gbp) of 100 random studies from the SRA. Construction time does not include the time of preprocessing the samples (graph cleaning). The parameters under the dashed line were estimated based on the key characteristics of the indexes shown above the dashed line. The best value for each characteristic is highlighted in bold.

| Characteristic | fast, RowFlat | fast, RowDisk | small, RowDisk | small, BRWT |
| --- | --- | --- | --- | --- |
| Graph (SuccinctDBG) | static | static | small | small |
| Annotation (RowDiff*$\ell$) | RowFlat | RowDisk | RowDisk | Multi-BRWT |
| Construction time, h | 24.0 | **23.6** | 23.8 | 24.8 |
| Size on disk $I$, GB | 51 | 49 | 37 | **32** |
| RAM usage $R_M(I)$, GB | 57 | 37 | **24** | 37 |
| Disk swap $R_L(I)$, GB | **0** | 25 | 25 | **0** |
| Search speed $S_{\text{search}}(I)$, Mbp/h | **713.1** | 562.2 | 284.6 | 144.9 |
| Alignment speed $S_{\text{align}}(I)$, Mbp/h | **24.5** | 20.8 | 8.3 | 6.3 |
| [dashed] Compression $c = D/I$, bp/byte | 188 | 195 | 259 | 299 |
| $r = R_M(I)/I$ | 1.12 | 0.76 | 0.65 | 1.16 |
| $l = R_L(I)/I$ | 0 | 0.51 | 0.68 | 0 |
| $s_{\text{search}}$, Mbp/h/GB | 14.0 | 11.5 | 7.7 | 4.5 |
| $s_{\text{align}}$, Mbp/h/GB | 0.48 | 0.42 | 0.22 | 0.20 |

representing the graph with SuccinctDBG (static) and the annotation with RowDiff<RowFlat> provides the highest query throughput. However, it required more RAM when querying compared to other representations. The index using SuccinctDBG (small) and RowDiff<Multi-BRWT>, on the contrary, achieves the best compression, but it provides lower throughput. Finally, the indexes representing the annotation with RowDiff<RowDisk> provide an excellent trade-off. They require significantly less RAM when querying by using additional space on SSD or NVMe SSD.

The k-mer search and sequence alignment throughput was measured by querying a batch of reads randomly sampled from the same data set. Namely, from those 5,184 samples, we randomly selected 300 and randomly picked 100 reads from each of them, which resulted in 30,000 reads of the total size 5.3 Mbp.

##### Final estimates for SRA

Finally, we used our methodology described in Section A.5.1 to estimate the cost of sequence search against the entire SRA with different hosting scenarios described in Section A.5.2. We parsed the list of all available compute instances of the current generation provided by the Amazon AWS provider in the US East (Ohio) region (data from https://calculator.aws/#/addService/ec2-enhancement/, accessed on 10.06.2023) and their prices for on-demand and dedicated (assuming prepayment for 3 years) use. We also extended this list with various compute instances provided by the Google Cloud Platform (data from https://cloud.google.com/products/calculator/, accessed on 10.06.2023).

For each instance type from the list and each version of MetaGraph, we estimated the number of machines of that type required to host the entire index of SRA using the parameters estimated in Section A.5.3. From this, we could immediately calculate the cost of serving the indexes on those machines (cost of the machines plus cost of storing the precomputed indexes) and the expected search throughput. For each hosting scenario and version of MetaGraph, we found the minimum hosting cost per year and then found an instance that would maximize the query throughput while still keeping the hosting cost at most 20% higher than the previously computed minimum. Note that since the hosting cost for 'MetaGraph on demand' only includes the cost of storing the preconstructed indexes, and hence, does not depend on the type of the compute instances used, the optimization is done to maximize the total query throughput, while for the other hosting scenarios

Table S-5: Costs for storing and searching against the SRA with MetaGraph in different representations ('MetaGraph on demand' uses the same index as 'MetaGraph fast, RowFlat') and Serratus. The search throughput was estimated for k-mer mapping. The effective query costs were computed for k-mer mapping (search) and alignment to labels (align). (\*) The indexing costs per year were estimated assuming that the SRA grows every year by the same absolute amount as in 2022. (†) The MetaGraph versions marked with † assume that the indexes are stored in fast local disk space and queried sequentially when a sufficiently large query batch is accumulated.

|  | Indexing cost |  | Hosting |  |  | Search | Query cost /Mbp |  |  |
| --- | --- | --- | --- | --- | --- | --- | --- | --- | --- |
| Method | before 2023 | then /year* | cost /year | node type | #nodes | Mbp/h | search | align |  |
| Storage of SRA | n/a | n/a | \$3,951,607 | AWS S3 buckets | n/a | 0 | n/a | n/a | |
| MetaGraph | fast, RowFlat | \$500,410 | \$133,213 | \$2,971,394 | x2gd.xlarge | 3373 | 2366 | \$0.14 | \$4.1 |
| | fast, RowDisk | \$499,883 | \$133,073 | \$1,940,876 | x2gd.xlarge | 2190 | 1211 | \$0.18 | \$4.9 |
| | small, RowDisk | \$500,226 | \$133,164 | \$1,264,034 | x2gd.2xlarge | 711 | 398 | \$0.35 | \$12.2 |
| | fast, RowFlat † | \$500,410 | \$133,213 | \$163,281 | i3en.3xlarge | 28 | 20 | \$0.71 | \$10.6 |
| | small, BRWT † | \$500,754 | \$133,304 | \$104,391 | i3en.3xlarge | 18 | 6.4 | \$1.44 | \$30.1 |
| | on demand | \$500,410 | \$133,213 | \$52,261 | e2-standard-32 | any | unlim. | \$0.10 | \$2.8 |
| Serratus | 0 | 0 | 0 | ((r5.c5).x/c5.)large | any | unlim. | n/a | \$1,736 | |

the optimization is done to primarily minimize the total yearly hosting cost. The results are shown in **Extended Data Table S-5**. (The detailed calculations with formulas can be found in the respective spreadsheet at https://github.com/ratschlab/metagraph_paper_resources.)

The indexing costs for each version of MetaGraph were estimated with the approach described in Section A.5.3 and include both the cost of preprocessing the samples as well as the index construction costs. While they make up a large figure in total, the indexing has to be done only once, and it takes only 2.2 cents when calculated per sample, which is roughly the price of storing this sample for 2 months with the Amazon AWS S3 service.

Once the indexes are constructed, they can be hosted in the cloud. The hosting costs, therefore, consist of two components: i) the cost of storing the indexes in the cloud and ii) the cost of running machines with the initialized indexes serving user queries. For the scenario 'MetaGraph on demand' (see **Extended Data Table S-5**), it is sufficient to only store the constructed indexes in fast common cloud storage to allow starting by the users any desired number of machines that would copy the indexes from the common storage and execute the requested queries, at their cost. Hence, the hosting (maintenance) costs would only include the cost of storing the index in that particular representation (SuccinctDBG (static) with RowDiff;RowFlat), which would be \$52,261 per year for the Amazon AWS S3 service. Note that due to the compression achieved during the indexing, storing the indexes is almost two orders of magnitude cheaper than storing the raw compressed samples, which is currently carried out by NCBI and ENA.

The other hosting scenarios ('fast, RowFlat', 'fast, RowDisk', etc.) assume not only storing the preconstructed indexes but constantly keeping them loaded in RAM of the compute machines. In this case, fast common storage such as AWS S3 is not required, and it is sufficient to store the indexes on regular HDD (e.g., EBS Cold HDD (sc1) Volumes provided at a price of \$0.015 per GB-month) to enable reloading the index, e.g., when a hosting machine unexpectedly shuts down. We calculated the storage costs for each representation of MetaGraph with the EBS Cold HDD (sc1) Volumes storage and added these to the hosting costs reported in **Extended Data Table S-5**. Besides those storage costs, the total yearly hosting costs we report include the cost of running the required number of compute machines of the selected type (see **Extended Data Table S-5**). To calculate these, we used the discounted pricing assuming a prepayment for 3 years of dedicated use. The effective query cost was calculated accordingly by dividing the hosting costs by the maximum throughput provided by a particular hosting scenario. The effective query costs for 'MetaGraph on demand' were calculated by dividing the price of machines of type e2-standard-

32 per time unit (provided by the Google Cloud Platform at the on-demand rate without discounts for committed use) by the estimated maximum query throughput it would achieve with the given number of virtual cores. Additionally, in this calculation, we did not assume that the indexes assigned to a specific machine for querying have to be simultaneously loaded in RAM since they can be queried sequentially with the same throughput when using multiple threads. As a result, the effective query costs for ‘MetaGraph on demand’ are estimated to be as low as \$2.8 per 1 Mbp of query sequences, which is over 600 times cheaper when compared to Serratus.

Last, one could argue that the indexing and hosting costs would grow exponentially with the exponential growth of the archive. However, when calculated per sample, these costs are not only fixed, but they are, in fact, three orders of magnitude lower than the actual cost of sequencing (the amortized cost of indexing and storing the index for a period of ten years is around 5 cents per sample while the sequencing currently costs around \$100 per sample and would unlikely be cheaper than 1\$ due to the required lab work and the cost of reagents). This suggests the feasibility of maintaining in the long term a system, which would allow cost-efficient sequence alignment against the entire SRA.

#### Chapter B

### Supplementary Figures

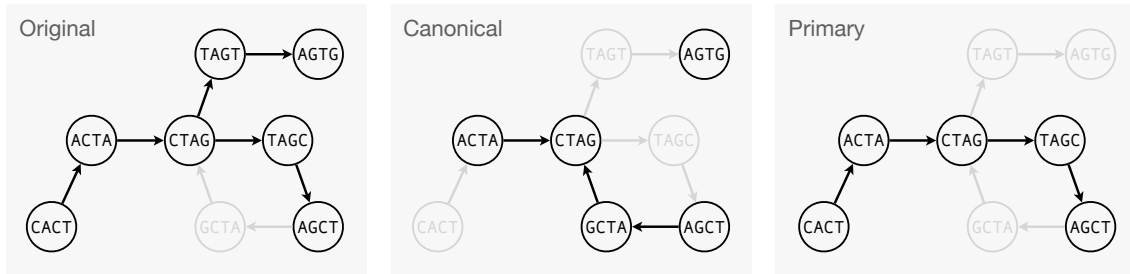

Figure S-5: Schematic illustration of canonical and primary graphs. **Left:** A De Bruijn graph of order $k = 4$ constructed from k-mers observed in the input sequences, e.g., CACTAGCT, CTAGTG. All k-mers (in this case only one k-mer GCTA) that did not actually occur in the input but could be present in the sample in reverse complement orientation are dimmed in gray. **Middle:** Canonical graph. All non-canonical k-mers are dimmed in gray and are represented implicitly by the explicitly stored canonical k-mers. For the BOSS table, however, all canonical and non-canonical k-mers are stored in this mode to reduce the number of disconnected components in the graph and thereby minimize the number of extra dummy k-mers. **Right:** Primary graph (i.e., a graph constructed from primary contigs). All non-primary k-mers are dimmed in gray.

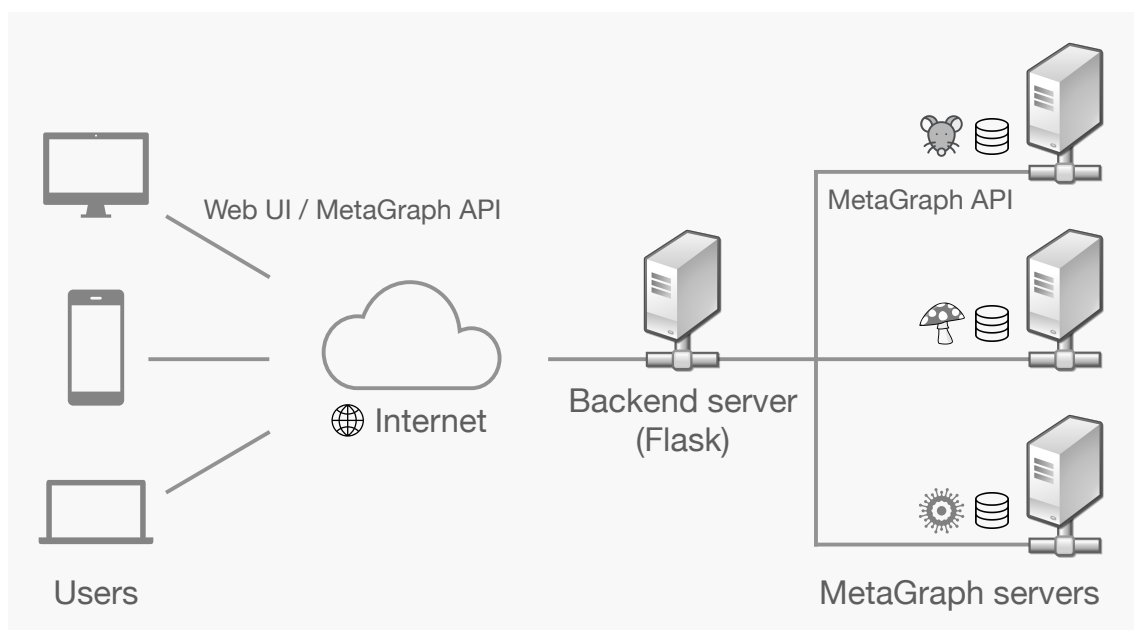

Figure S-6: Architecture of MetaGraph Online — The client-server architecture of MetaGraph Online. The backend server (middle) generates dynamic web pages and transforms user queries to search requests sent to the remote servers hosting MetaGraph indexes. It also provides an API equivalent to that of the MetaGraph server by forwarding requests sent to specific endpoints to their respective MetaGraph servers.

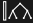

Home Search Align Graphs

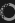

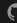

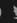

BioRxiv

##### MetaGraph: Search DNA Sequences

```
TTTCACTCTTTGATAGCAGATGCTTAGTACTAACTAAGTCCTCAAGATTGTGAGTCAGTCCTTCATTCTTCTACTGATAGTACTAGTATGACTGATCTCCCG
CTGCACUTAAACCAAAAGATACACTACTTAATTACCACTAGAAATATACAAATCATGCGATCATAGAAATCGAGACAACTTTTCCCAAGCAGGGTTT
```

Select graph:
SRA-Fungi

Minimum k-mer matches: 100%

☐ Search with alignment

Search SRA-Fungi

##### Search results

Download as csv

Show 10 entries

Search:

| # | sample | k-mer matches |
| --- | --- | --- |
| 1 | ERR1885701 | 180 |
| 2 | ERR1885702 | 180 |
| 3 | ERR1885703 | 180 |
| 4 | ERR1885704 | 180 |
| 5 | ERR1885705 | 180 |
| 6 | ERR1885706 | 180 |
| 7 | SRR5449081 | 180 |
| 8 | SRR2841901 | 180 |
| 9 | SRR4026169 | 180 |
| 10 | ERR6829475 | 180 |

Showing 1 to 10 of 96 entries

Previous 1 2 3 4 5 ... 10 Next

Figure S-7: MetaGraph Online — Web user interface of the MetaGraph Online search engine for sequence search.

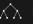

Home Search Align Graphs

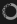

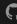

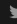

BioRxiv

##### List of Graphs

| Index | Input | k | Num k-mers | Num labels | Indexing mode | API endpoint |
| --- | --- | --- | --- | --- | --- | --- |
| GnomAD (release 3.0) | N/A | 41 | 29,015,017,946 | 29 | ordinary | /api/gnomad |
| Kingsford (2,652 RNA-Seq) | 8.0 Tbp | 21 | 10,165,997,234 | 2,586 | with k-mer counts | /api/kingsford |
| MetaSUB (k=41) | 7.2 Tbp | 41 | 166,983,379,394 | 4,220 | ordinary | /api/metastub41 |
| RefSeq (85k taxID; release 97; with coordinates) | 1.7 Tbp | 31 | 626,753,663,468 | 85,375 | with k-mer positions | /api/refseq85_coord |
| SRA-Fungi | 160 Tbp | 31 | 264,681,928,748 | 121,900 | ordinary | /api/sra_fungi |
| SRA-Metazoa | 156 Tbp | 31 | 750,598,764,890 | 241,384 | ordinary | /api/sra_metazoa |
| SRA-Metazoa (1,000 studies) | 119 Tbp | 31 | 625,622,500,650 | 67,390 | ordinary | /api/sra_metazoa1k |
| SRA-Microbe | 221 Tbp | 31 | 101,733,896,434 | 446,506 | ordinary | /api/sra_microbe |
| SRA-Mouse (Mus Muculus) | 147 Tbp | 31 | 139,978,792,470 | 57,938 | ordinary | /api/sra_mus_muculus |
| Tara Oceans (genomes with coordinates) | 61.9 Gbp | 31 | 26,511,636,608 | 34,815 | with k-mer positions | /api/tara_genomes_coord |
| Tara Oceans (scaffolds) | 357 Gbp | 31 | 121,058,940,696 | 318,205,057 | ordinary | /api/tara_assemblies |
| UHGG All | 710 Gbp | 31 | 33,120,094,297 | 286,997 | ordinary | /api/uhgg_all |
| UHGG Catalogue | 11.2 Gbp | 31 | 9,676,446,464 | 4,644 | ordinary | /api/uhgg |

Figure S-8: List of indexes available on MetaGraph Online — Web view of indexes hosted on the MetaGraph Online search engine for sequence search.
